# Exploring Microbial Diversity and Functional Potential along the Bay of Bengal Coastline in Bangladesh: Insights from Amplicon Sequencing and Shotgun Metagenomics

**DOI:** 10.1101/2023.04.26.538428

**Authors:** Salma Akter, M. Shaminur Rahman, Hazrat Ali, Benjamin Minch, Kaniz Mehzabin, Md. Moradul Siddique, Syed Md. Galib, Farida Yesmin, Nafisa Azmuda, Nihad Adnan, Nur A Hasan, Sabita Rezwana Rahman, Mohammad Moniruzzaman, Md Firoz Ahmed

## Abstract

Although the Bay of Bengal (BoB) is the world’s largest bay, possessing distinct physiochemical properties, it has garnered little research focus concerning its microbial diversity and ecological importance. Here, we present amplicon (16S and 18S) profiling and shotgun metagenomics data regarding microbial communities from BoB’s eastern coast, viz., Saint Martin and Cox’s Bazar, Bangladesh. From the 16S sequencing data, Proteobacteria appeared to be the dominant phylum in both locations, with *Alteromonas*, *Methylophaga*, *Anaerospora*, *Marivita*, and *Vibrio* dominating in Cox’s Bazar and *Pseudoalteromonas*, *Nautella*, *Marinomonas*, *Vibrio*, and *Alteromonas* dominating the Saint Martin site. From the 18S sequencing data, Ochrophyta, Chlorophyta, and Protalveolata appeared among the most abundant eukaryotic divisions in both locations, with significantly higher abundance of Choanoflagellida, Florideophycidae, and Dinoflagellata in Cox’s Bazar. Functional annotations revealed that the microbial communities in these samples harbor genes for biofilm formation, quorum sensing, xenobiotics degradation, antimicrobial resistance, and a variety of other processes. Together, these results provide the first molecular insight into the functional and phylogenetic diversity of microbes along the BoB coast of Bangladesh and lay the foundation for further in-depth assessment of microbial community dynamics and functional potential in the context of global change in this region.

## 1. Introduction

The oceans cover 70% of the earth’s surface and are home a myriad of microorganisms, all of which contribute to the survival of life on earth ^1^. These microorganisms are important for the health of aquatic ecosystems that vary geographically due to environmental conditions, community adaptability, and anthropogenic impacts ^2,3^. Global change is expected to influence both the mean and variance of environmental parameters in the open sea, with global pH decreases and ocean surface water temperature rises ^4,5^. As microbes play a significant role in marine nutrient cycling, climate models should account for changes in microbial community structure and biogeochemical activities ^6–8^.

The coastline of the Bengal delta comprises the Bay of Bengal (BoB), the largest bay in the world ^9^. Due to considerable influence by seasonal natural disasters such as monsoon rainfalls, climate disasters, and human development, the BoB gets a significant flux of fresh and cold river water into this semi-enclosed tropical ocean basin in the northeast Indian Ocean ^9,10^. Rising surface water temperatures in the BoB have led to heightened stratification in the water column, creating zones characterized by depleted oxygen and nutrient levels ^10^.

The coastal ecosystem provides vast scope for economic development through the establishment of ports, fisheries industries, gas fields, oil refineries, and naval stations. Despite enormous economic contributions to coastal countries like Bangladesh, India, Myanmar, and Sri Lanka, the BoB ecosystem is extremely underexplored. Several reports from neighboring countries showed investigative outcomes on oceanography, phytoplanktonic diversity, and stratification-induced nutrient cycling, but without a notable focus on microbial composition through advanced molecular studies ^9,11–13^.

Multiple studies have found that BoB oceanic characteristics have a significant impact on the composition and metabolic diversity of the marine microbiome ^14,15^. Recent large-scale projects in conjunction with modern DNA sequencing technologies have made significant contributions to the microbial characterization of numerous marine ecosystems, ranging from the Arctic Ocean to the tropics ^16–19^. Several studies from India have reported the microbial diversity of the surface and sub-surface regions of BoB ^13,20,21^, but no study has been performed in Bangladesh yet, despite the substantial economic and ecological importance of BoB to Bangladesh. These coastal regions of Bangladesh play an important economic role because they are the most visited tourist destination in the country^22,23^ and the largest source of fisheries-based rural markets, supplying a significant portion of the country’s fish^24^.

The present study aims to identify the prokaryotic and eukaryotic diversity of the microbiome of two distinct coastal regions of Bangladesh – Cox’s Bazar and Saint Martin. We performed 16S and 18S high-throughput amplicon sequencing to determine the diversity of prokaryotic and eukaryotic microbes. We then conducted concordant shotgun metagenomic sequencing to assess several key functional aspects of the community. Specifically, we sought to investigate the prevalence of pathogenic microbes and traces of antimicrobial resistance, which are strong indicators of anthropogenic disturbances in marine ecosystems ^25^.

## 2. Methodology

### 2.1 Sample collection

The seawater samples were collected in duplicates from two distinct coastal regions of Bangladesh: Cox’s Bazar and the Saint Martin. The sampling was done on March 2 and 3, 2022 during low tide. The samples were collected in 1 L sterile sampling bottles at 1.5-meter depth from the surface water. The bottles were sealed underwater and transported to the Microbiology laboratory at Jahangirnagar University, Savar, Dhaka, Bangladesh for further processing. The samples from Saint Martin were labeled as S5, S6, S7 and S8 and the samples from Cox’s Bazar were labeled as S9, S10, S11 and S12. The geographical location of each sampling sites is available in Supplementary Data-1. Water samples from each site were taken in sterile beaker and physicochemical parameters like temperature, pH, salinity, and total dissolved solids (TDS) were measured using suitable handheld devices (Hanna, USA).

### 2.2 Total DNA extraction from water samples/ Molecular Processing

The water samples were initially passed through Whatman filter paper no. 1 (pore size 11µm) to get rid of any large debris. The water filtrates were then filtered through the Millipore filtration unit, firstly through 0.45 µm membrane and subsequently through 0.20 µm membrane. The filtrate was discarded, and the filter papers were folded in 5 ml sterile tubes and stored at −80°C for DNA extraction. From these filter papers sample DNA was extracted using DNeasy PowerWater Kit (QIAGEN) according to the manufacturer’s protocol. The purified DNA extracts from duplicates samples of a single site were combined together and were quantified to determine concentration and relative purities, prior to sending for 16S and 18S rDNA based metagenomic sequencing done by EzBiome, USA. For whole genome metagenomic (shotgun) sequencing, equal quantity of the extracted DNA from both 0.45 and 0.20 µm membranes from representative four sampling sites of two locations were combined as pooled samples (Cox’s Bazar (S2) and Saint Martin (S1)).

### 2.4 Library Preparation and Sequencing

The amplification of prokaryotic DNA was achieved by targeting the V3–V4 region of 16S rRNA gene with 30 μL final volume containing 15 μL of 2 × master mix (BioLabs, USA), 3 μL of template DNA, 1.5 μL of each V3–V4 forward and reverse primers, 341F (5′-CCT ACG GGNGGCWGCAG-3′) and 806R (5′-GACTACHVGGGTATCTAATCC-3′), respectively^26^. The remaining 9 μL of DEPC treated ddH2O. A 25 cycle of PCR amplification including initial denaturation at 95 °C for 3 min, denaturation at 95 °C for 30 s, primer annealing at 55 °C for 30 s and elongation at 72 °C for 30s was performed for bacterial DNA with the final extension of 5 min at 72 °C in a thermal cycler (Analytik Jena, Germany).

To amplify DNA, the universal eukaryotic primers set 1391F (5ʹ-GTA CAC ACC GCC CGTC-3ʹ) / EukBr (5ʹ-TGA TCC TTC TGC AGG TTC ACC TAC-3ʹ) spanning the V9 region of 18S rRNA gene were utilized^26^. PCR mixture for the amplification of fungal DNA was the same as the one used for prokaryotic DNA. For eukaryotic DNA, a thirty-five cycles of PCR amplification were run with the temperature profile of initial denaturation at 94 °C for 3 min, denaturation at 94 °C for 45 s, annealing at 57 °C for 1 min, elongation at 72 °C for 1.5 min and final extension of 10 min at 72 °C. After electrophoresis, the PCR amplicons were visualized in 1.5% agarose gel prepared in 1 × TAE buffer. Agencourt Ampure XP beads (Beckman Coulter, Brea, USA) were used for PCR products purification, and the Nextera XT index kit (Illumina, San Diego, USA) for paired-end library preparation according to Illumina standard protocol (Part# 15,044,223 Rev. B). Paired-end (2 × 300 bp reads) sequencing of the prepared library pools was performed using MiSeq high throughput kit (v3 kit, 600 cycles) with an Illumina MiSeq platform (Illumina, USA)^27,28^.

### 2.5 Bioinformatics data processing

The generated FASTQ files were evaluated for quality using FastQC v0.11^29^. Adapter sequences, and low-quality ends per read were trimmed by using Trimmomatic v0.39 with a sliding window size of 4; a minimum average quality score of 20; minimum read length of 40 bp^30^. After quality control, there were an average of 9305 pairs of reads for 16S samples (minimum = 7476 and maximum = 11961 pairs) and an average of 34,144 pairs of reads for 18S samples (minimum = 51681 and maximum = 22392 pairs). QIIME 2 (2022.2), an integrated pipeline was used for OUT clustering, phylogenetic estimation and taxonomic assignment^31^. VSEARCH metagenomics algorithm integrated in QIIME 2 was employed for read joining, dereplicate-sequences, *de novo* clustering (OUT clustering with 99 % identity), *de novo* chimera checking (exclude chimeras and “borderline chimeras”) ^32^. To generate a tree for phylogenetic diversity analyses, MAFFT ^33^ was used for alignment and FastTree (v2.1.8) was used to build the tree ^34^.

For taxonomic assignment, Greengenes (v13_5) database (99% OTU and taxonomy) used for prokaryotic taxonomic assignment (16S) and SILVA (v132_99) database (99% OTU and taxonomy) were also used for eukaryotic taxonomic assignment^35,36^. The reference database was trained using the 16S and 18S sequencing primer pairs using a naive-bayes classifier^37,38^. Classify-sklearn algorithms were utilized to classify the assigned OTU for prokaryotic and eukaryotic samples^39,40^.

### 2.6 Statistical analysis

The downstream analysis, which included alpha and beta diversity, microbiological composition, and statistical comparison, were performed using the Phyloseq (version 4.2) package ^41,42^ for R (v 4.2.1) ^43,44^. Observed, Chao1, Shannon, Simpson, InvSimpson, and Fisher alpha diversity were estimated and plotted by using “Vegan”, “ggplot2”, and “ggpubr” R packages. The Wilcoxon sum rank test in the “microbiomeutilities” R package (https://microsud.github.io/microbiomeutilities/) was used to evaluate the differences in microbial diversity and abundance between two locations. Beta diversity was measured with the principal coordinate analysis (PCoA) using Bray–Curtis, weighted unifrac, and unweighted unifrac dissimilarity matrices, and permutational multivariate analysis of variance (PERMANOVA) with 999 permutations was used to estimate a p-value for differences between two locations. The non-metric multidimensional scaling (NMDS) method was also applied for the above-mentioned distance metrics including PERMANOVA. Phyloseq, Vegan, microbiome utilities, and ggplot2 packages were employed for taxonomic comparison and plotting ^41,45–48^. To analyze and illustrate the data, the R packages Hmisc and corrplot were used ^49–51^.

### 2.7 Shotgun metagenomic sequencing, and sequence reads preprocessing

Both Cox’s Bazar and Saint Martin’s samples were combined into two different pools before submission to shotgun metagenomic sequencing. Shotgun metagenomic (WMS) libraries were prepared with Nextera XT DNA Library Preparation Kit and paired-end (2 × 150 bp) sequencing was performed on a NovaSeq 6000 sequencer (Illumina Inc., USA) from EzBiome, USA. The generated FASTQ files were evaluated for quality using FastQC v0.11^29^. Adapter sequences, and low-quality ends per read were trimmed by using Trimmomatic v0.39 with a sliding window size of 4; a minimum average quality score of 20; minimum read length of 50 bp^30^. In the end, the trimmed read counts for S1 and S2 were 33.94 and 31.8 million, or 92.20 and 92.37% of the total raw read counts, respectively.

### 2.8 Taxonomic mapping, classification, and phylogenetics study

CZID (previously IDseq), a real time microbiome characterization pipeline (v7.1) ^52^ and EzBioCloud taxonomic profiling ^53^ were used for taxonomic identification of the short read sequences. CZID is an open-source cloud-based pipeline for taxonomic assignments against the NCBI non-redundant (NR) database with NRL (NRL; non-redundant nucleotide alignment length in bp) ≥ 50 and NR % identity ≥ 80. CZID applies host filtering, alignment with minimap2 ^54^ assembly with SPAdes ^55^ and blast for taxonomic assignment.

Bacteria, Archaea, Virus and cdf (https://www.ncbi.nlm.nih.gov/refseq/) were also added to the Kraken2 database ^56^. After acquiring a list of candidate species, a custom bowtie2^57,58^ database was built utilizing the core genes and genomes from the species found during the first step. The raw sample was then mapped against the bowtie2 database using the very sensitive option and a quality threshold of phred33. Samtools^59,60^ was used to convert and sort the output BAM file. Coverage of the mapped reads against the bam file was obtained using Bedtools^61,62^. Then, to avoid false positives, using an in-house script, we quantified all the reads that mapped to a given species only if the total coverage of their core genes (archaea, bacteria) or genome (fungi, virus) was at least 25%. Finally, species abundance was calculated using the total number of reads counted and normalized species abundance was calculated by using the total length of all their references.

### 2.9 Shotgun Sequence Assembly

Short reads from both metagenomic libraries were quality trimmed using Trim Galore (https://github.com/FelixKrueger/TrimGalore) with default parameters^63^. The trimmed data was assembled using metaSPADES^64–66^ with default parameters and a minimum contig size of 1500 base pairs. Gene prediction of the metagenomic contigs was done using Prodigal^67^ with the meta option.

### 2.10 Functional Profiling and BRITE Hierarchy Analysis

For each sample, functional annotations were obtained by matching each read against the KEGG database using DIAMOND ^54,68–70^. DIAMOND was executed using the blastx parameter, which converts each metagenomic read into multiple amino acid sequences by generating all six open reading frame variations, and then matches it against the pre-built KEGG database. After quantifying all the KEGG orthologs present, minpath^71^ was used to predict the presence of KEGG functional pathways. The KEGG BRITE database is a collection of BRITE hierarchy files, called htext (hierarchical text) files, with additional files for binary relations. The htext file is manually created with in-house software called KegHierEditor^72^. The htext file contains “A”, “B”, “C”, etc. at the first column to indicate the hierarchy level, and may contain multiple tab-delimited columns. Thus, the htext file is like an Excel file with the additional first field for the hierarchy level. The BRITE hierarchy file has been created to represent the functional hierarchy of KEGG objects identified by the KEGG Identifiers.

### 2.11 Antimicrobial Resistance Genes (ARGs) and Virulence Factor-associated Genes (VFGs) Profiling

For antimicrobial resistance profiling, two different pipelines were used. The first is the AMR++ pipeline with the Microbial Ecology Group (MEG) antimicrobial resistance database (MEGARes v3.0.0)^73–75^. The short reads were aligned to the MEGARes database using Burrows-Wheeler Aligner (BWA)^76^, with the gene fraction (the percentage of genes that were matched to by at least one sequencing read) set to ≥80%. Contigs obtained from CZID pipeline and refined bins were also aligned against the MEGARes database with ≥80% identity and ≥80% subject coverage. In addition, the EzBioCloud pipeline was also used to assign ARGs from short reads. Antibiotic resistance gene profiles were produced by using a pre-built bowtie2^57^ database composed of NCBI’s National Database of Antibiotic Resistant Organisms (NDARO, (www.ncbi.nlm.nih.gov/pathogens/antimicrobial-resistance/) reference genes. Each read of the metagenome sample was mapped against these genes using bowtie2 with the very-sensitive option, and the output was then converted and sorted by Samtools^60^. Finally, for each gene found, depth and coverage were calculated by using Samtool’s mpileup script. We used the same pipelines mentioned above to find virulence factor-associated genes (VFGs) from the Virulence Factors of Pathogenic Bacteria (VFDB) database^74,77–79^.

## 3. Results

### 3.1 Physicochemical properties of water samples

A total of 8 samples were collected from the coastal regions of Cox’s bazar and Saint Martin Bangladesh during the 2nd and 3rd of March 2022. (Fig. 1A). The samples from Cox’s Bazar had an average pH of 7.3, while the ones from Saint Martin’s had a slightly higher average pH of 7.425. The maximum salinity, TDS, and temperature in samples from Cox’s Bazar were, 35 units (average = 32.75), 7028 units (average = 6656.5), and 30.7 °C (average = 28.03 °C) respectively, and in samples from Saint Martin were 36 units (mean = 35.75), 7580 units (mean = 6593.25), and 30.7 °C (mean = 28.03 °C) respectively (Fig. 1B). No statistically significant variations have been observed in the physicochemical parameters among samples from these two locations (t-test, p > 0.05) (Fig. 1B) (Supplementary Data 1).

**Fig. 1:**
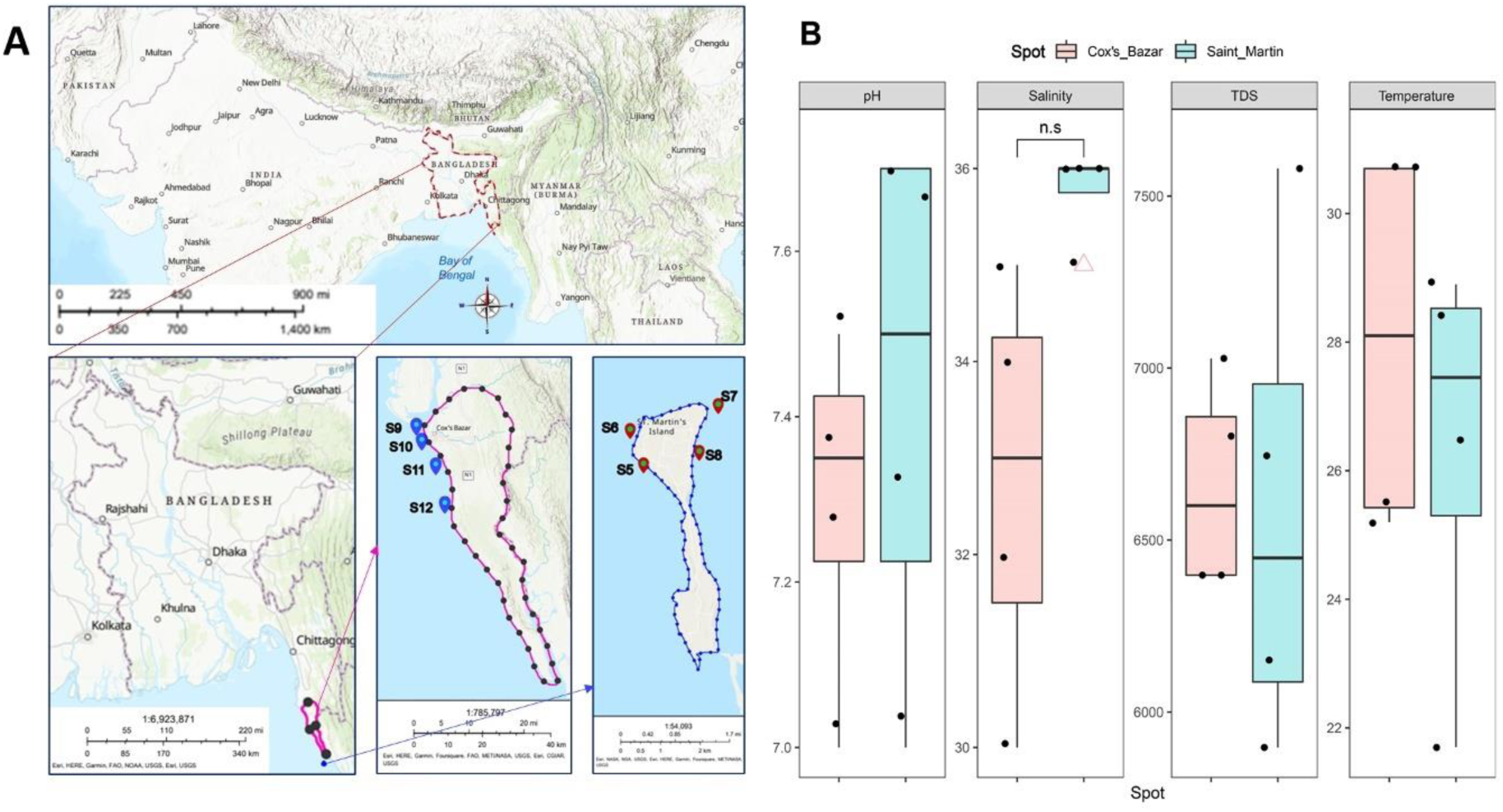
Sampling location and their physicochemical properties. (A) Two sampling locations (Cox’s Bazar and Saint Martin) are indicated (yellow rectangle). (B) The physicochemical parameters (pH, salinity, TDS and Temperature) of each are plotted on boxplots and comparisons were made with t-test. The map was constructed using ArcGIS online platform.

### 3.2 16S and 18S Microbiome diversity

#### 3.2.1 Bacterial and Archaeal Diversity from 16S amplicons

We were able to get a total of 397 OTUs (Operational Taxonomic Units) from the 16S microbiome sequences derived from V3-V4 amplicons of all the samples. After clustering and filtering for chimeras, the Observed, Chao1, Shannon, Simpson, InvSimpson, and Fisher indices were examined for within-sample diversity (Alpha diversity), but the results showed that there was no significant difference (Wilcoxon signed-rank test, p > 0.05) between the two locations in terms of bacterial and archaeal diversity (Fig. 2A). Principal coordinate analysis (PCoA) with Bray Curtis distance (Fig. 2B), weighted unifrac distance (Fig. 2C), and unweighted unifrac distance (Fig. 2D) showed that there were no significant differences between the two sampling locations (beta diversity) (PERMANOVA, p > 0.05). Similar results were obtained using the non-metric multidimensional scaling (NMDS) technique, with no discernible differences (PERMANOVA, p > 0.05) (Fig. 2E-G).

**Fig. 2:**
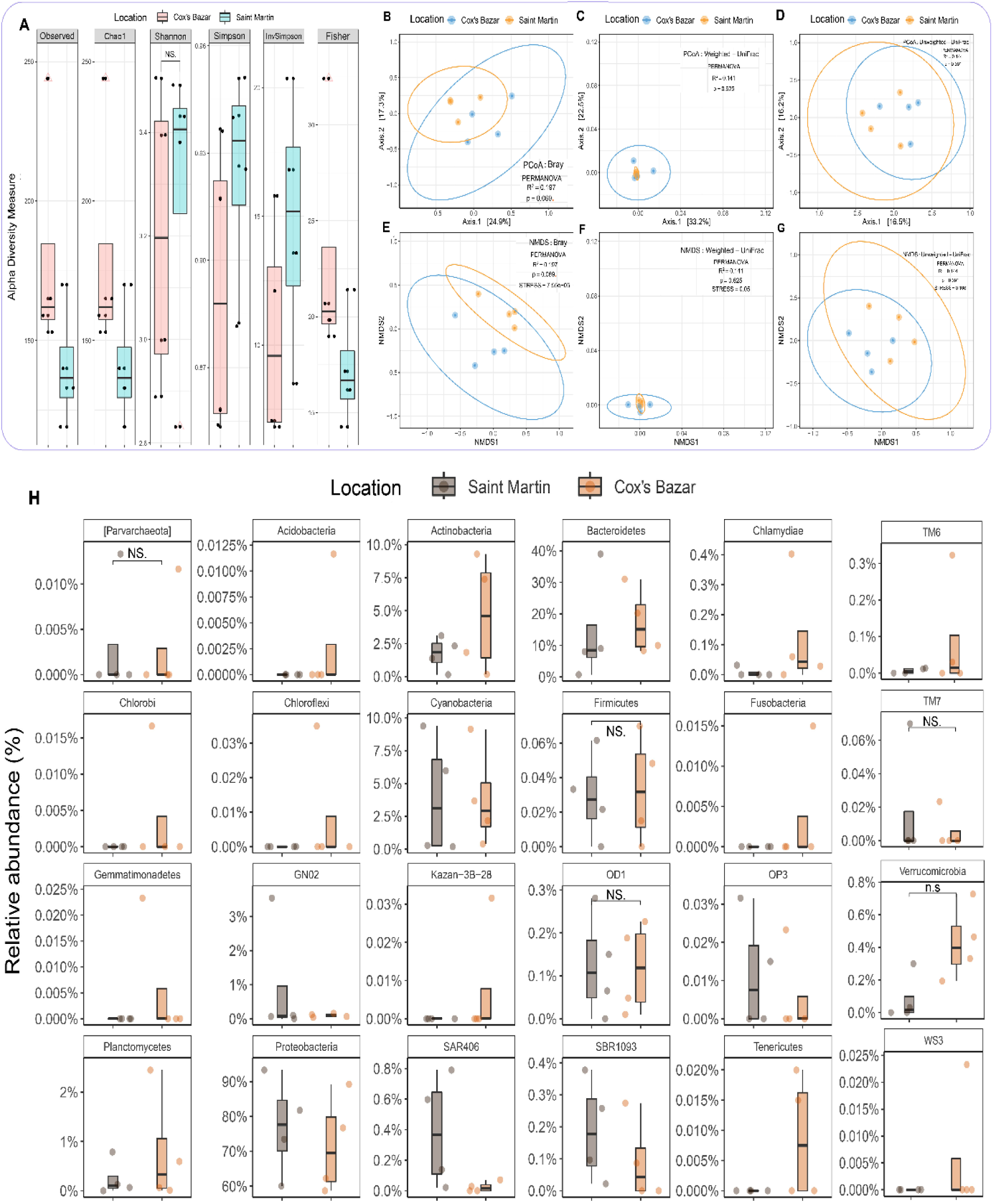
Prokaryotic and eukaryotic microbial alpha- and beta-diversity based on 16S and 18S taxonomic abundance. **(A)** For the prokaryotic (bacteria and archaea) microbial community of Cox’s Bazar and Saint Martin samples, the observed species, Chao1, Shannon, Simpson, InvSimpson, and Fisher diversity (Alpha diversity) indices were estimated. X-axis represents the location and y-axis represents the alpha diversity measure. The diversity for each is plotted using boxplots, and the pairwise Wilcoxon sum rank test is used to compare them. (B-G) Beta diversity measures of the prokaryotic (bacteria and archaea) microbial community. Principal coordinate analysis (PCoA) **(B-D)** and non-metric multidimensional scaling **(E-G)** were performed using Bray, Weighted-Unifrac, and Unweighted-Unifrac distance metrics for the two locations of samples. Permutational multivariate analysis of variance (PERMANOVA) was performed with 999 permutations to estimate a significance (p-value) for differences between two locations. PERMANOVA with 999 permutations was used to determine the significance (p-value) of differences between two locations. Significance level (p-value) 0.0001, 0.001, 0.01, 0.05, and 0.1 are represented by the symbols “****”, “***”, “**”, “*”, and “n.s”, respectively. Stress value represents the goodness of fit of NMDS (> 0.2 Poor, 0.1-0.2, Fair, 0.05-0.1 Good, and <0.05 Excellent). **(H)** Comparison of relative abundance of twenty-five prokaryotic phyla in the two different locations (Cox’s Bazar and Saint Martin). The diversity for each phylum is plotted on boxplots and comparisons are made with Wilcoxon sum rank test.

Our study revealed the presence of a total of 24 bacterial phyla and one archaeal phylum (Parvarachaeota) in the sequence data. 16 bacterial phyla were found in the Saint Martin region, in contrast to the 24 that were found in Cox’s Bazar (Supplementary Data-3: Figure-1). All 16 phyla that were found in Saint Martin were also found in Cox’s Bazar. More than 98% of the bacterial phyla in the Cox’s Bazar area were comprised of Proteobacteria (71.7%), Bacteroidetes (17.4%), Actinobacteria (4.7%), Cyanobacteria (3.8%), and Planctomycetes (0.8%). On the other hand, almost 97% of all phyla in the Saint Martin were Proteobacteria (77.1%), Bacteroidetes (14.2%), Cyanobacteria (3.95%), and Actinobacteria (1.7%) (Figure-2H).

From the QIIME2 analysis of 397 OTUs, a total of 138 bacterial genera were identified from both locations, among them 120 and 88 genera were found in Cox’s Bazar and Saint Martin respectively. Notably, the top ten genera from Cox’s Bazar had 84.4% relative abundance, consisting of sequences that could not be assigned to any known phyla (47.21%), *Alteromonas* (10.27%), *Methylophaga* (8.57%), *Anaerospora* (6.31%), *Marivita* (2.89%), *Vibrio* (1.97%), *Synechococcus* (1.85%), *Sediminicola* (1.79%), *Nautella* (1.78%), and *Pelagibacter* (1.75%). On the contrary, the top ten genera from Saint Martin had 94.2% relative abundance, consisting of sequences with unknown assignment (40.72%), *Pseudoalteromonas* (9.39%), *Nautella* (6.96%), *Marinomonas* (6.92%), *Vibrio* (5.64%), *Alteromonas* (4.85%), *Synechococcus* (3.49%), *Polaribacter* (3.23%), *Candidatus Portiera* (2.72%) and *Pelagibacter* (2.26%) (Supplementary Data 1). Cox’s Bazar had significantly higher abundance for *Antarctobacter* (Wilcoxon rank test p-value = 0.029), *Formosa* (p-value = 0.029) and *Marivita* (p-value = 0.021) and Saint Martin had significantly higher abundance for *Oleibacter* (p-value = 0.029), and *Rhodovulum* (p-value = 0.029) (Supplementary Data-3: Figure-2).

#### 3.2.2 Diversity of microbial eukaryotes from 18S amplicons

After clustering and screening for chimeras from the V9-amplicons of 18S microbiome sequencing, we were able to get a total of 693 OTUs (Operational Taxonomic Units) from all samples (S5-S12). Observed, Chao1, Shannon, Simpson, InvSimpson, and Fisher indices no significant difference within sample (alpha) diversity (Wilcoxon signed-rank test, p >0.05) between the Cox’s Bazar and Saint Martin’s samples. (Fig. 3A). Principal coordinate analysis (PCoA) using Bray Curtis distance (Fig. 3B), weighted unifrac distance (Fig. 3C), and unweighted unifrac distance (Fig. 3D) revealed considerable differences (PERMANOVA, p < 0.05) between the two sampling locations of samples. An NMDS approach revealed the same significant difference (PERMANOVA, p < 0.05) (Fig. 3E-G). A closer examination revealed that three and nineteen divisions were unique to Saint Martin and Cox’s Bazar, respectively, with the remaining 22 divisions shared by both locations (Supplementary Data-3: Fig. 1B). In both Cox’s Bazar and Saint Martin, a large proportion of the OTUs could not be assigned to known divisions (74.27% and 88.85% respectively). In Cox’s Bazar, the most abundant divisions found are Ochrophyta (11.44%), Chlorophyta (4.72%), Fungi (1.98%), Labyrinthulomycetes (1.76%), Protalveolata (1.61%), Cercozoa (1.19%), and Choanoflagellida (1.15%). In the Saint Martin samples, Chlorophyta (7.77%) was the most abundant flowed by Protalveolata (1.88%), Ochrophyta (0.49%) and Fungi (0.37%) (Supplementary Data 1). Between the two sites, only Choanoflagellida (p=0.021), Florideophycidae (p=0.021), and Dinoflagellata (p=0.029) were found to have significantly different abundance, all being higher in Cox’s bazar (Figure-3H).

**Fig. 3:**
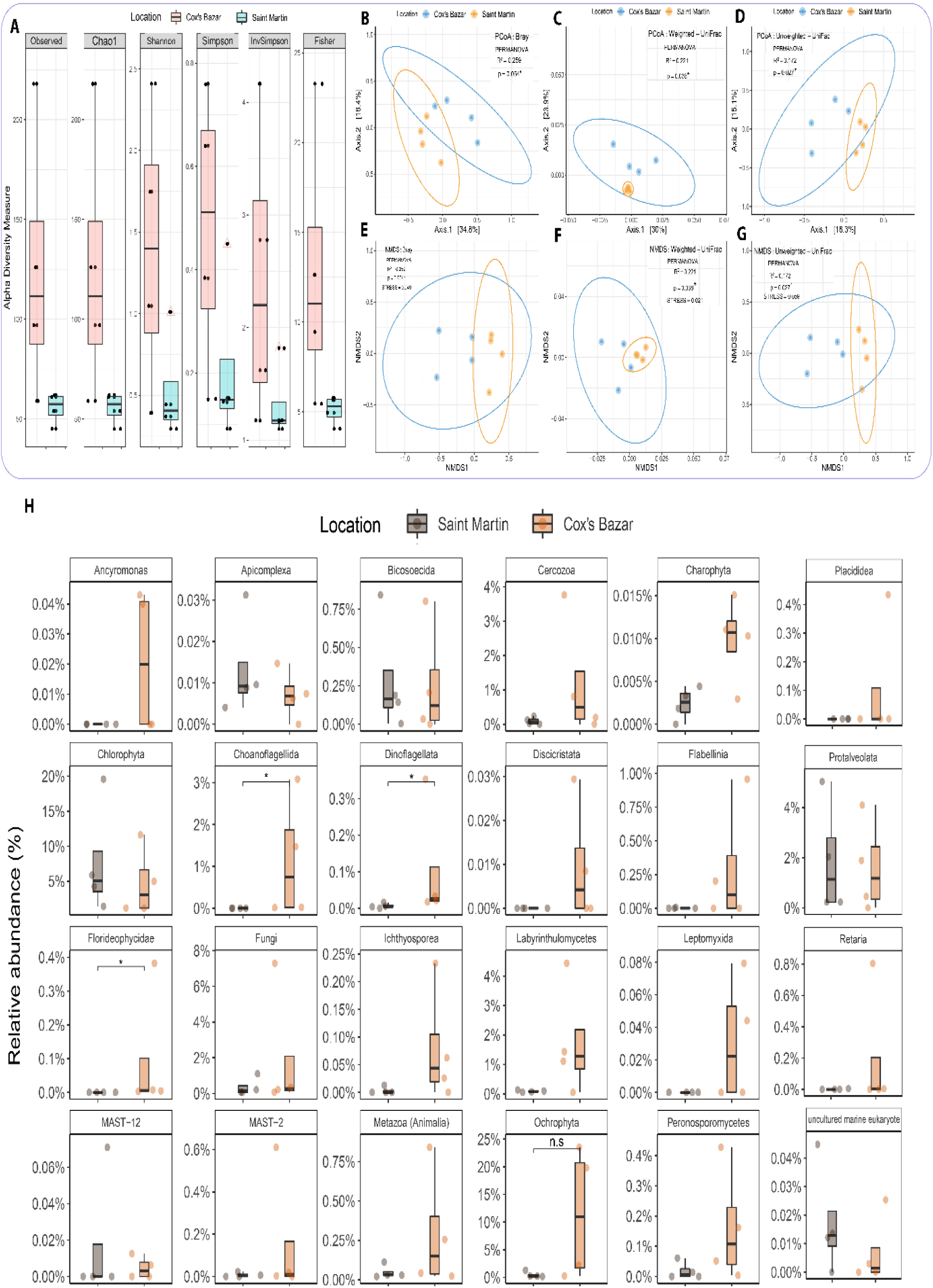
**(A)** The observed species, Chao1, Shannon, Simpson, InvSimpson, and Fisher diversity (Alpha diversity) measures were used to estimate the Eukaryotic microbial community diversity of Cox’s Bazar and Saint Martin samples as described for the prokaryotic microbes. **(B-G)** Beta diversity of the eukaryotic microbial community was estimated here as described in Figure-2 (B-G). **(H)** Comparison of relative abundance of twenty-five eukaryotic divisions in the two different locations (Cox’s Bazar and Saint Martin). The diversity for each division is plotted and differences were tested using Wilcoxon sum rank test. Significance level (p-value) 0.0001, 0.001, 0.01, 0.05, and 0.1 are represented by the symbols “****”, “***”, “**”, “*”, and “n.s”, respectively.

#### 3.2.3 Site specific relative abundance of different genera

The relative abundance of the dominant genera in the samples of eight sites showed significant variations in the dominance of bacterial genus (Figure-4A). Among the top 20 genera, *Alteromonas* appeared to be the most dominant one with highest abundance in S9 sample, followed by *Pseudoalteromonas* which was most abundant in S6. The next abundant genera, *Anaerospora,* was dominant in S11. Among the other genus *Methylophaga* and *Polaribacter* mostly belonged to S12 and S5 respectively. Other genera like *Vibrio* and *Nautella* were distributed in all the samples.

**Figure 4:**
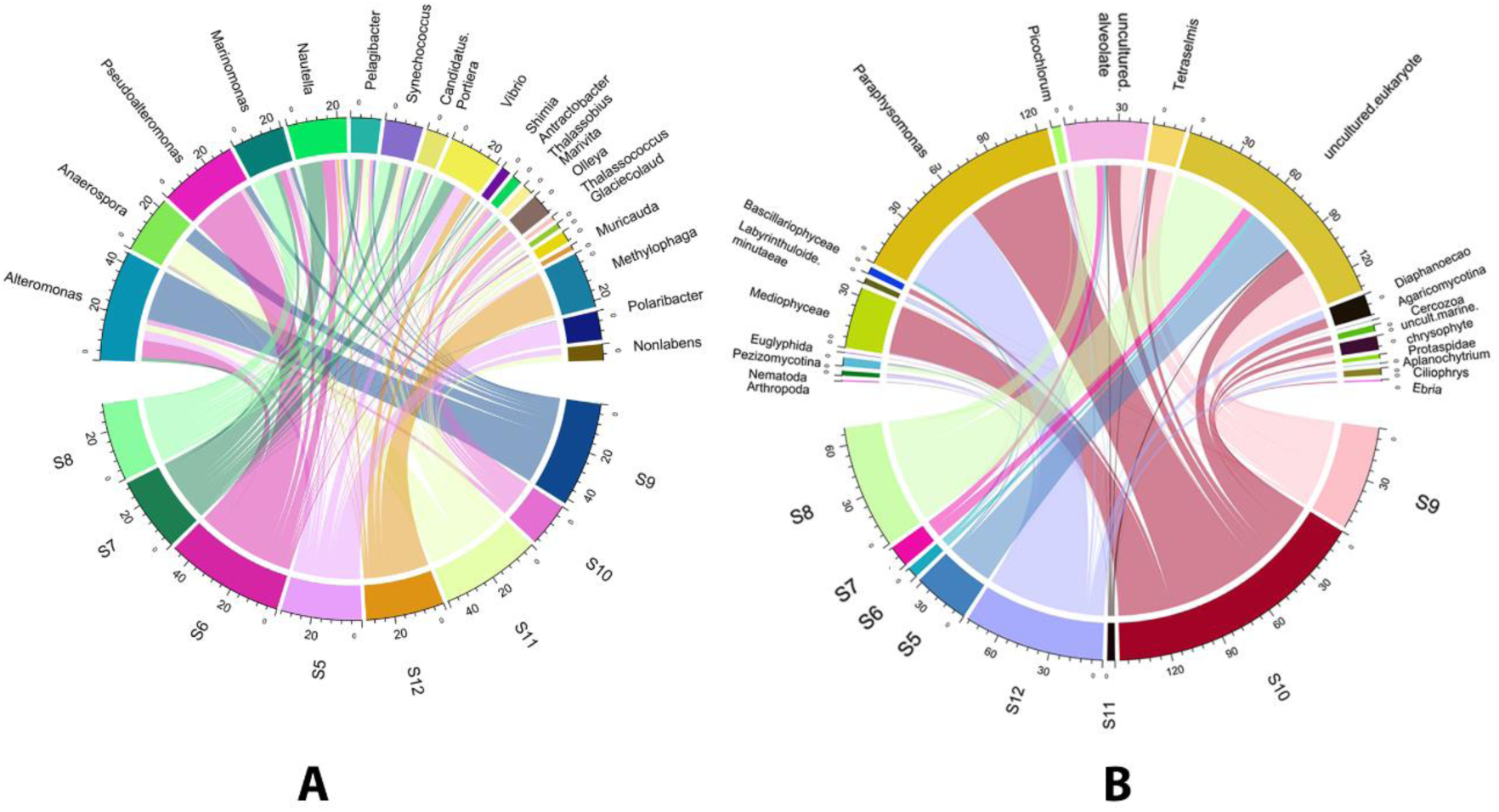
Circos representation of relative abundance for the top 20 prokaryotic genera (A) from 16S rRNA sequence data and top 20 eukaryotic genera (B) from 18S rRNA rRNA sequence data obtained across different sampling sites. Sample S5-S8 belong to Saint Martin and S9-S12 belong to Cox’s Bazar. The representing values are the 1st percentile of the actual read numbers.

The 18S sequence data showed the maximum relative abundance read for S10 and that was followed by S12, S8 and S9 (Figure-4B). Among sites, the majority of taxa remained unknown. *Paraphysomonas* was the most abundant genera and was almost equally distributed to S10 and S12 sites. *Mediophyceae*, the next dominant eukaryotic genera were found exclusively in S10. Another most abundant taxa, uncultured alveolates, was mostly associated to S9 and S8 however, but were also present in other samples. Overall, the differences in relative abundance for the top 20 genera was more noticeable for the eukaryotic organisms than prokaryotic ones in the sampling sites.

### 3.3 Impact of environmental conditions on microbial community composition

The influences of physicochemical factors on the relative abundance of prokaryotic and eukaryotic microbial communities of the samples revealed that Parcubacteria (also known as Candidate Phylum OD1 bacteria (OD1)) showed significant negative correlation with pH (Spearman correlation; r > −0.86, p < 0.01). Planctomycetes demonstrated a substantial positive association with TDS (Spearman correlation; r > 0.78, p < 0.01) and a significant negative correlation with temperature (Spearman correlation; r > −0.78, p < 0.01) (Figure-5A). Fungi and Ichthyosporea showed strong negative correlation with pH (Spearman correlation; r > - 0.86, p < 0.01) and salinity (Spearman correlation; r > −0.87, p < 0.01) respectively (Figure - 5B).

**Figure-5: Pairwise.**
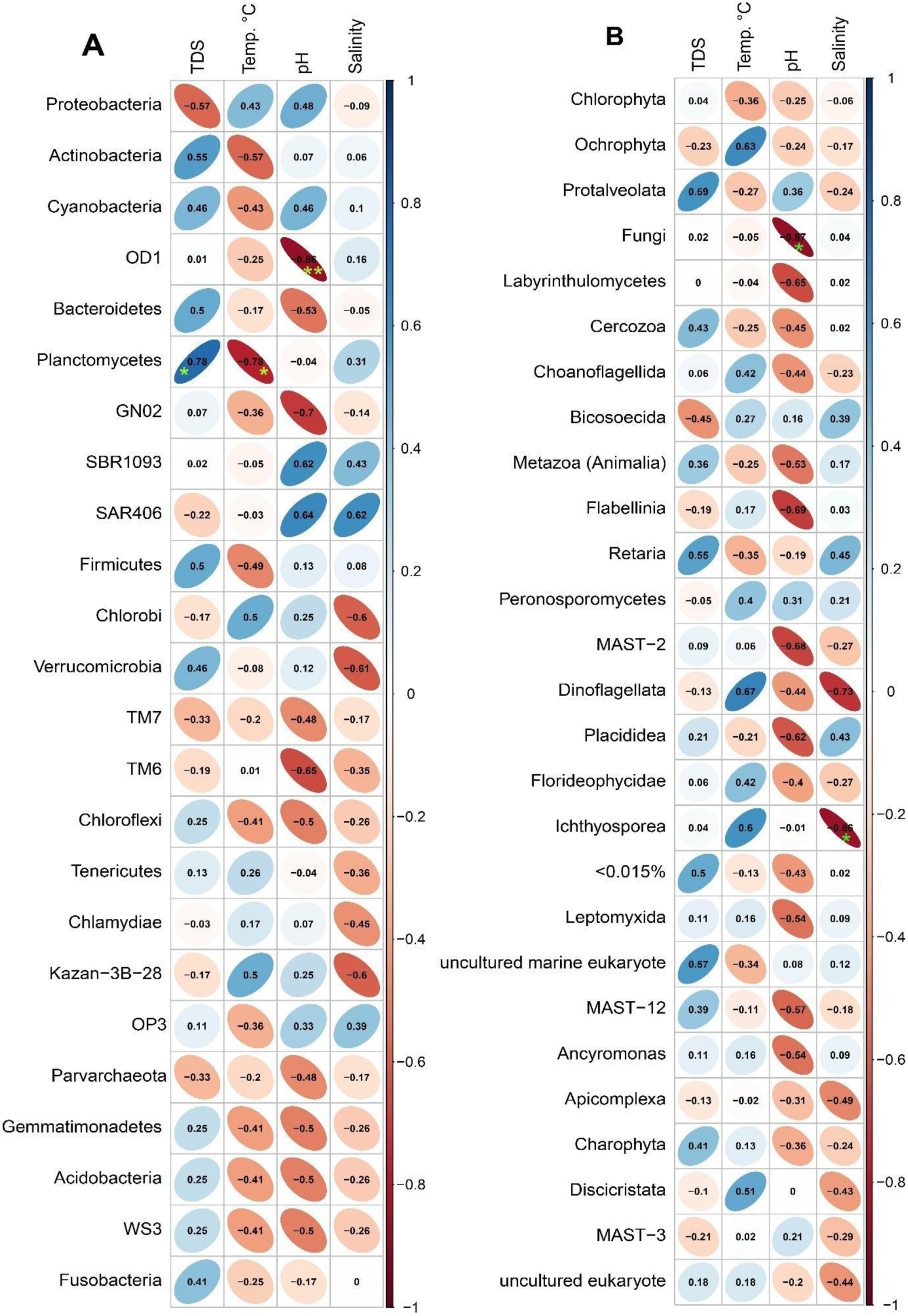
Spearman’s correlation of physicochemical parameters and microbial phyla (prokaryotic) and division (eukaryotic) level. (A) Correlation with physicochemical parameters (TDS, temperature, pH, and salinity) with 24 phyla of prokaryotes detected in the study areas. (B) Correlation with physicochemical parameters with top 26 divisions (> 0.015%) of eukaryotes detected in the study areas. The numbers represent the Spearman’s correlation coefficient (r). Blue and red indicate positive and negative correlations, respectively. The color density, ellipse size, and numbers reflect the scale of correlation. *Significance level (*p < 0.05; **p < 0.01; ***p < 0.001).

### 3.4 Shotgun Metagenomic Analysis

#### 3.4.1 Taxonomic Composition of Prokaryotic and Eukaryotic Microbial Community

For assessment of overall community composition and relative functional profiling of surface microbiome of the two coastal regions of BoB, we also performed shotgun metagenomic sequencing using the pooled DNA samples (S1=Saint Martin; S2= Cox’s bazar). From the taxonomic profiling data, both S1 and S2 sample showed to harbor bacteria, eukaryotes, archaea and viruses (Supplementary Data-2). Among them, 99.13% and 99.33% sequences revealed presence of bacteria in S1 and S2 respectively, followed by eukaryotes (0.01%, 0.03%), viruses (0.80%, 0.64%) and archaea (0.02%, 0.01%).

*Altermonas* appeared as the most prevalent bacterial genera in both locations, followed by *Methylophaga* for Cox’s Bazar and *Vibrio* for Saint Martin (Figure-6). Among the other genera *Pseudoalteromonas*, *Rhodobacteraceae*, *Cognatishimia*, *Marinomonas*, *Phaeobacter*, and *Proteobacter* are fairly abundant in from both locations. At the species level, *Alteromonas macleodii* was predominant in both locations followed by *Methylophaga aminisulfidivorans*, *Alteromonas* sp., *Rhodobacteraceae bacterium*, *Methylophaga sulfidovorans*, *Donghicola tyrosinivorans*, *Alteromonas abrolhosensis*, and *Rhodobacteraceae* bacterium for Cox’s Bazar, and *Alteromonas macleodii*, *Methylophaga aminisulfidivorans*, *Alteromonas* sp., *Rhodobacter*aceae bacterium, *Methylophaga sulfidovorans*, *Donghicola tyrosinivorans*, *Pseudoalteromonas phenolica*, *Alteromonas abrolhosensis*, *Rhodobacteraceae bacterium*, *Vibrio natriegens*, *Cognatishimia maritima* and *Cognatishimia active* for Saint Martin.

**Figure 6:**
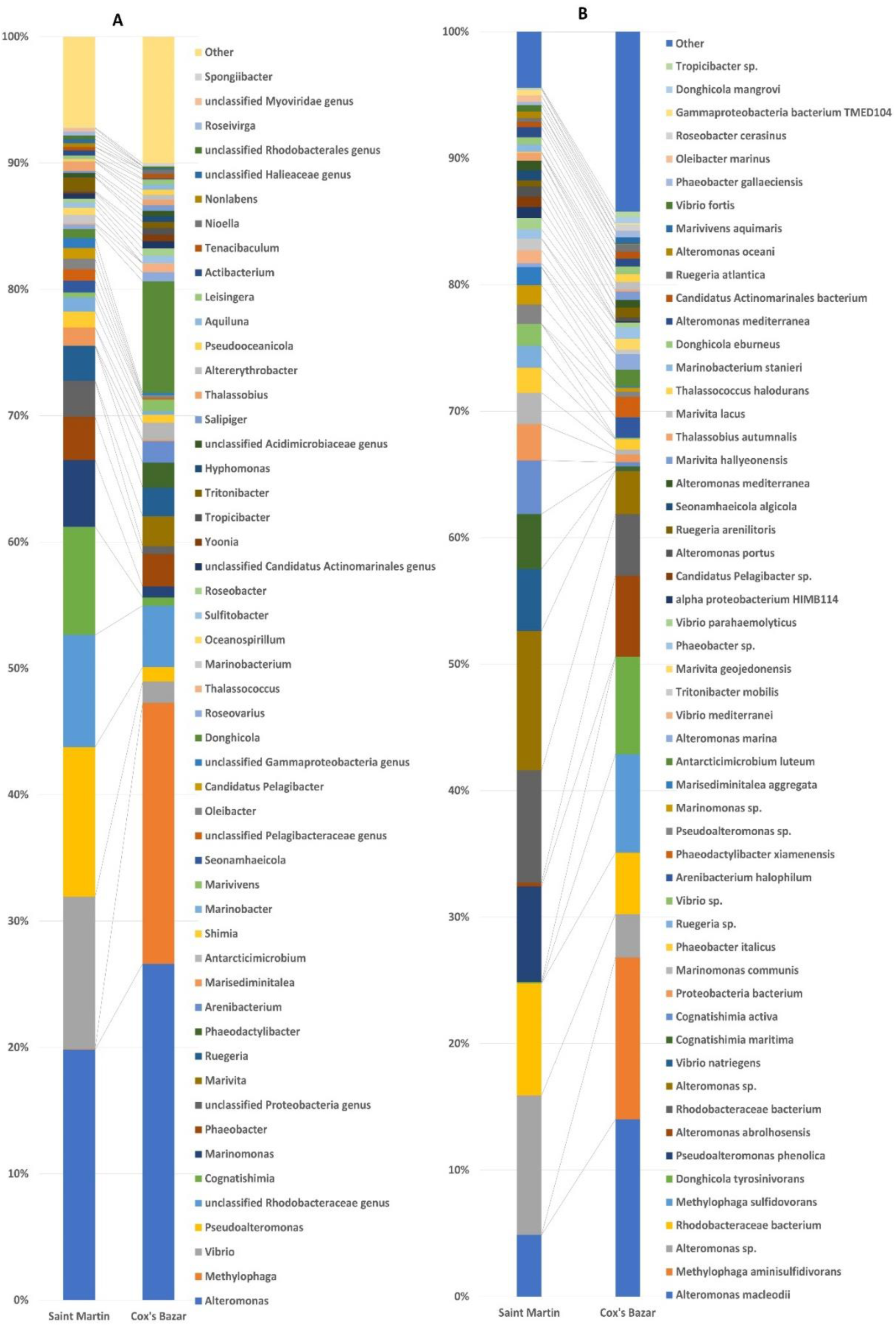
The (A)genera and (B)species level taxonomic profile of microbes obtained from shotgun metagenomic sequencing of Saint Martin (S1) and Cox’s bazar (S2) samples. Stacked bar plots showing the relative abundance and distribution of the top 50 genus and species. The distribution and relative abundance of the microbes in the study metagenomes are also available in Supplementary Data-2.

**Figure-7:**
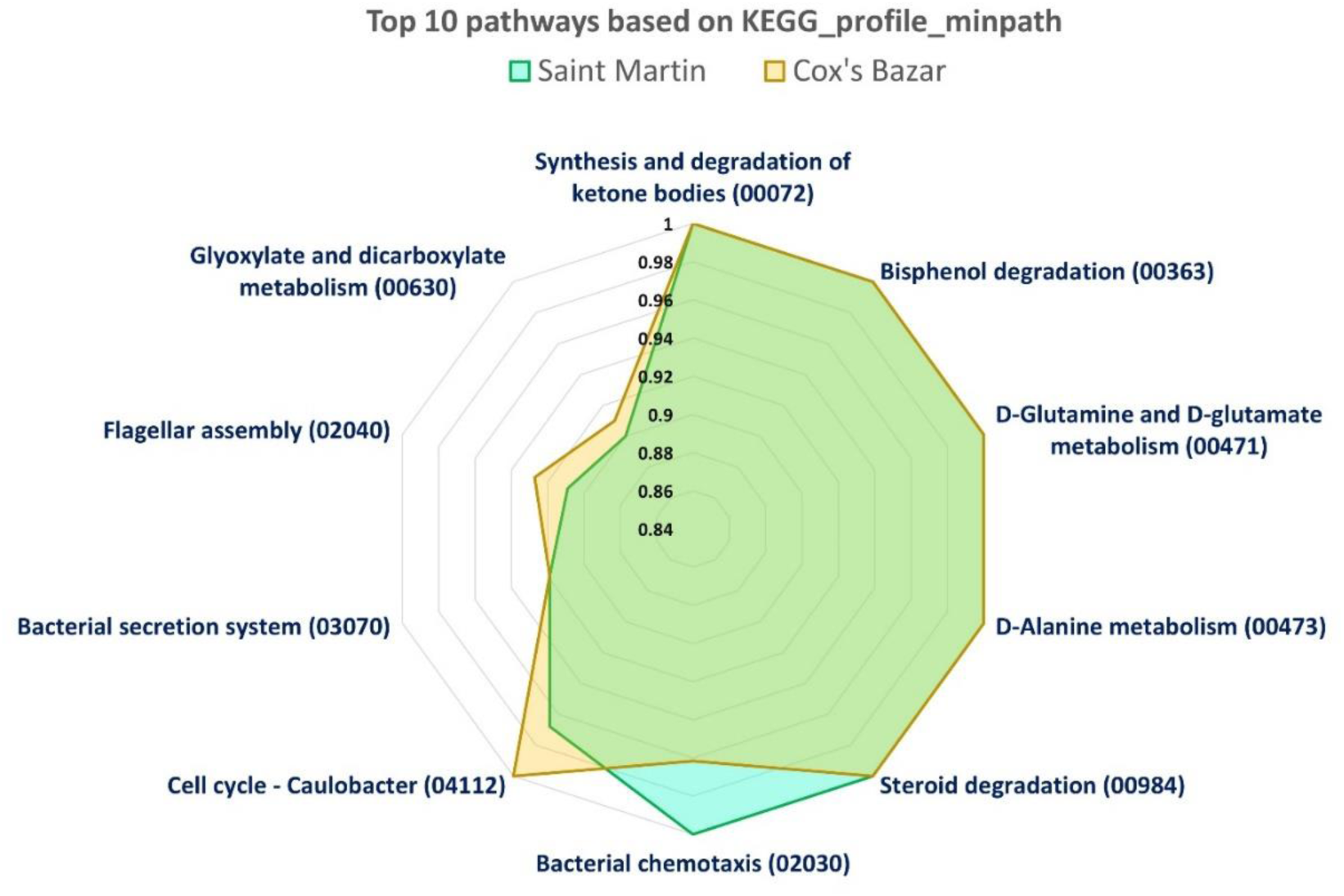
Most abundant (Top 10) pathways present in with the marine microbiome in BoB, Bangladesh (based on KEGG_profile_minpath).

#### 3.4.2 Functional Profiling of BoB Microbiome

All levels of functional gene profiling using KEGG Orthology (https://www.genome.jp/kegg/ko.html) revealed differential abundance of metabolic genes in two samples. The most abundant metabolic category was the BRITE Hierarchies category (KO09180) present in Cox’s bazar and St martins with a relative abundance of 0.3789 and 0.3826, respectively. Notably, the KEGG Orthology derived functional gene identification showed the presence of human disease-causing genes in the both samples.

The top 15 BRITE level B found in Cox’s Bazar and Saint Martin were Protein families involved in signaling and cellular processes (ko09183), genetic information processing (ko09182), amino acid metabolism (ko09105), carbohydrate metabolism (ko09101), metabolism (ko09181), metabolism of cofactors and vitamins (ko09108). It is interesting to note that the distribution of BRITE level B categories is similar between the two locations, with only small differences in the abundance of each category (Supplementary data-3: Table-1).

In BRITE level C functional gene annotation by KEGG-Orthology revealed that the two sources of marine water samples have similar relative abundances of proteins. For example, both locations have relatively high levels of transporters, enzymes with EC (Enzyme Commissioner) numbers, DNA repair and recombination proteins, and transfer RNA biogenesis proteins. There were also some differences between the two sources. Cox’s Bazar has higher relative abundances of glycine, serine, and threonine metabolism proteins, as well as porphyrin metabolism proteins, while Saint Martin has higher relative abundances of ABC transporters and peptidases and inhibitors. The most abundant KEGG orthologous group in both locations is K02014 (TC.FEV.OM), which is involved in the transport of amino acids, indicating a higher demand for amino acids in these locations, possibly due to high metabolic activity or protein synthesis. The second most abundant orthologous group in Saint Martin is K03406 (mcp), which is involved in bacterial chemotaxis, whereas in Cox’s Bazar K20276 (bapA) is the second highest, which is involved in the formation of biofilms. This suggests that bacterial motility may be important in Saint Martin, while biofilm formation is more important in Cox’s Bazar. Figure-6 illustrates the relative abundance of top 10 metabolic genes prevalent in the functional microbiome of two samples, determined from shotgun metagenome sequences of S1 and S2. Importantly, the relative abundance of Bis-phenol degradation metabolism is higher in both samples, indicating the presence of potential microbial communities capable for possible photodegradation of bisphenol-A (BPA) which is a harmful component found in hard plastics, water bottles etc. The abundance of D-glutamate and D-Glutamine metabolism indicates the continuous fixation of atmospheric nitrogen by the marine bacteria and anabolic utilization of these amino acids for biosynthesis of proteins, nucleic acids in microorganisms.

#### 3.4.3 Antibiotics resistance gene families prevalent in coastal water microbiome of Saint Martin and Cox’s Bazar

In total, 54 antimicrobial and metal resistance genes (Supplementary Data-3: Table 3,4) were detected in the coastal water samples from BoB considering the gene coverage above 80%. Among them, 17 and 48 genes belong to S1 and S2, respectively. Only 11 AMR genes were found in both samples, whereas 6 and 37 genes were unique to S1 and S2 sample respectively. Saint Martin (S1) sample had relatively a smaller number of resistance genes where macrolide-resistance being the most abundant one, followed by aminoglycoside-resistance and quinolone-resistance. On the other hand, Cox’s Bazar (S2) samples had nearly three times more resistance than S1 samples with phenicol resistance gene being the most abundant one, followed by resistance to tetracycline, quinolone, macrolide and sulfonamide. Cox’s Bazar samples also encoded genes for resistance to various biocides and metals (Table-1). No resistance genes for tetracycline, phenicol and sulfonamides with >80% gene coverage have been found in S1 samples. Likewise, resistance genes for trimethoprim (with >80% coverage) have not been identified in S2 samples.

**Table 1:**
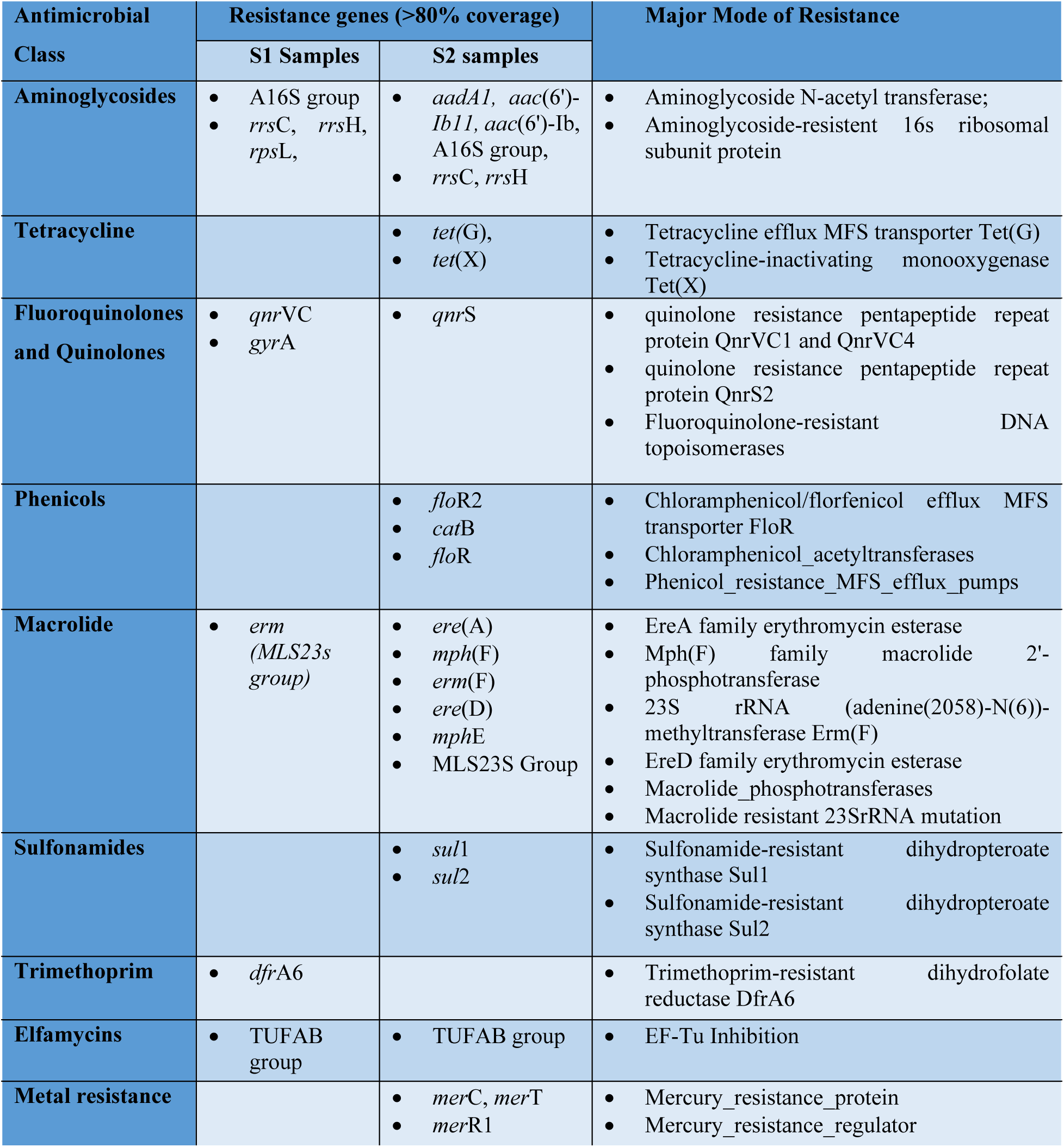
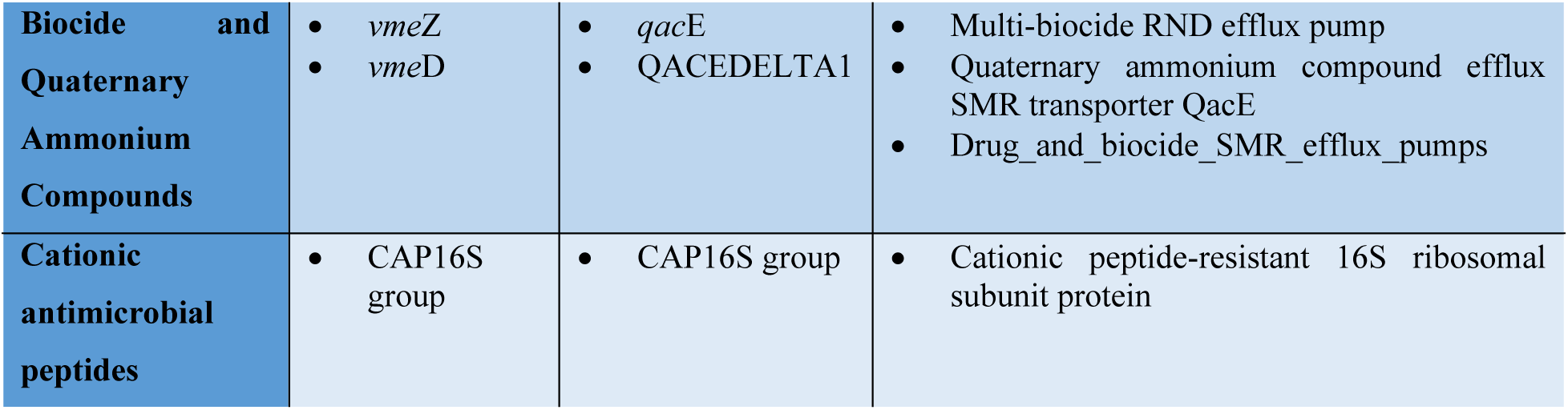
Antimicrobial resistance gene profiling for S1 and S2 samples.

| Antimicrobial Class | Resistance genes (>80% coverage) |  | Major Mode of Resistance |
| --- | --- | --- | --- |
|  | S1 Samples | S2 samples |  |
| **Aminoglycosides** | A16S group<br>*rrsC*, *rrsH*, *rpsL*, | *aadA1*, *aac(6')-Ib11*, *aac(6')-Ib*, A16S group,<br>*rrsC*, *rrsH* | Aminoglycoside N-acetyl transferase;<br>Aminoglycoside-resistant 16s ribosomal subunit protein |
| **Tetracycline** |  | *tet(G)*,<br>*tet(X)* | Tetracycline efflux MFS transporter Tet(G)<br>Tetracycline-inactivating monooxygenase Tet(X) |
| **Fluoroquinolones and Quinolones** | *qnrVC*<br>*gyrA* | *qnrS* | quinolone resistance pentapeptide repeat protein QnrVC1 and QnrVC4<br>quinolone resistance pentapeptide repeat protein QnrS2<br>Fluoroquinolone-resistant DNA topoisomerases |
| **Phenicol** |  | *floR2*<br>*catB*<br>*floR* | Chloramphenicol/florfenicol efflux MFS transporter FloR<br>Chloramphenicol_acetyltransferases<br>Phenicol_resistance_MFS_efflux_pumps |
| **Macrolide** | *erm* (*MLS23s* group) | *ere(A)*<br>*mph(F)*<br>*erm(F)*<br>*ere(D)*<br>*mphE*<br>MLS23S Group | EreA family erythromycin esterase<br>Mph(F) family macrolide 2'-phosphotransferase<br>23S rRNA (adenine(2058)-N(6))-methyltransferase Erm(F)<br>EreD family erythromycin esterase<br>Macrolide_phosphotransferases<br>Macrolide resistant 23SrRNA mutation |
| **Sulfonamides** |  | *sul1*<br>*sul2* | Sulfonamide-resistant dihydropteroate synthase Sul1<br>Sulfonamide-resistant dihydropteroate synthase Sul2 |
| **Trimethoprim** | *dfrA6* |  | Trimethoprim-resistant dihydrofolate reductase DfrA6 |
| **Elfamycins** | TUFAB group | TUFAB group | EF-Tu Inhibition |
| **Metal resistance** |  | *merC*, *merT*<br>*merR1* | Mercury_resistance_protein<br>Mercury_resistance_regulator |
| **Biocide and Quaternary Ammonium Compounds** | • *vmeZ*<br>• *vmeD* | • *qacE*<br>• QACEDELTA1 | • Multi-biocide RND efflux pump<br>• Quaternary ammonium compound efflux SMR transporter QacE<br>• Drug_and_biocide_SMR_efflux_pumps |
| **Cationic antimicrobial peptides** | • CAP16S group | • CAP16S group | • Cationic peptide-resistant 16S ribosomal subunit protein |

#### 3.4.4 Virulence factor associated gene families prevalent in coastal water microbiome of Saint Martin and Cox’s Bazar

From the analysis of functional properties of the prevalent microbiome of BoB, several genes related to virulence factors have been identified. The EzBioCloud and AMR++ pipelines both identified bacterial pathogenic genes mostly related to flagellar motility, such as *flgB, flgC, flgD, mshA, fliA* etc (Supplementary Dat-3: Table-5). Other genes for chemotaxis (*cheY*), transport protein (*pyuC, pysC*) and type II secreteion system protein (*epsE, epsG*) have been identified, which are involved in flagellar motility, nutritional uptake of metal Fe-like metal ions and secretion of effector moieties for flagella formation. Interestingly, most of the virulence genes identified from S1 sample had gene coverage >80%, whereas no genes from S2 samples had above 80%. Regardless of the coverage, shotgun metagenome sequence analysis of both samples has been determined to have significant presence of virulence genes which indicate that the coastal water of both locations is harboring pathogenic organisms. Notably, taxonomic identifications revealed presence of a number of pathogenic bacteria in the samples, justifying the source of virulence genes.

## 4. Discussion

Coastal microbiome research, particularly in the context of Bangladesh’s south and south-east coast, is still in its infancy. As a part of the Indian Ocean, the third largest oceanic division of the world ^80^, and being surrounded by three different countries, BoB provides ecological habitats and niches for an enormous diversity of microbial groups ^13,81^. A recent study conducted by Ghosh *et al* (2022) revealed that the bacterioplankton community in multiple locations of BoB showed the dominance of Proteobacteria, Bacteroidetes, and Firmicutes as well as the nitrogen-fixing groups such as Nitrospirae, Lentisphaerae, Chloroflexi, and Planctomycetes ^82^. Another metagenome-based study of deep-sea sediment samples from 3000m depth of BoB revealed the dominance of Proteobacteria followed by Bacteroidetes, Firmicutes, Cyanobacteria and Actinobacteria ^13^. The BoB possesses a large oxygen minimum zone (OMZ) which causes the shifting of microbial and planktonic communities due to the continuous variation of ocean water conditions ^83–86^. A study conducted by Bowie Gu ^87^ revealed the simultaneous shift of microbial community and functional profiles along with the oxygen concentrations and an evident role of *Trichodesmium* bloom in carbon and nitroigen availability resulting in OMZ formation in BoB. Several recent studies have explored the microbial and phytoplankton composition of different zones of BoB using next generation sequencing methods to identify the pools of microbiomes in culture-independent manner ^88^. Angelova et al. revealed that the diversity of planktonic microbial communities varies with vertical differentiation of population regardless of the sampling locations ^14^. To understand the diversity and functional potential of marine microbial communities and factors that influence their community dynamics, cutting-edge technologies like high-throughput metagenomic and meta-transcriptomic sequencing are widely used.

In this study, we investigated the microbial profile of two distinct coastal sites of Bangladesh. Our two study areas are around 50 nautical miles away and the samples from Cox’s Bazar and Saint Martin did not vary significantly from one another in terms of the tested physicochemical parameters. Bacteria are abundant and prevalent in marine ecosystems, playing a vital role in biogeochemical cycles and can account for up to 70% of total biomass in surface ^89^ and 75% in deep waters ^90,91^. However, their diversity and composition are frequently affected by a range of environmental conditions. In our study from shotgun sequencing data, we found more than 99% of the sequences belonged to bacterial kingdom and that was followed by viruses for both locations. Our sampling approach likely allowed for higher proportion of planktonic bacteria to be captured - as we passed the water samples through filters of pore size 11µm, which excluded some nanoplanktons (2-20 µm) and all microplanktons (20-200 µm). In addition, it might have also excluded microbial communities in association with particles and/or forming biofilms. Subsequently, the samples were passed through 0.45 µm followed by 0.22 µm membranes. The later approach removed many of the Femtoplanktons (0.01–0.2 µm) i.e., viruses. Therefore, only the cell associated viruses and viruses larger than 0.2 µm were contained in the membranes, making our samples contain mostly the picoplankton (0.2–2 µm) such as bacteria and archaea, and some viruses. Additionally, deeper sequencing and higher sample volume would potentially lead to a better estimate of the microbial diversity in our samples. Regardless of these limitations, our shotgun, 16S and 18S metagenomic sequencing revealed presence of at least 60 different phyla, total of 397 prokaryotic OTUs representing 24 bacterial phyla and one archaeal phylum, and 693 OTUs for eukaryotes representing 44 divisions.

In a recent study of deep sea sediment from BoB, 19 phyla were identified using a nanopore based approach ^13^. Interestingly, another 16S amplicon-based study from BoB’s oxygen minimum zones (OMZs) and non-OMZs on the Indian coasts identified over 4000 OTUs with more than 70% reads assigned to bacterial and 30% reads to archaeal domains. This massive difference in the OTU number could be due to differences in the utilization of reference databases, OTU threshold (99% vs 97% identity), and differences in sampling sites and zones.

The vast majority of eukaryotic OTUs from Cox’s Bazar (74.27%) and Saint Martin (88.85%) could not be assigned to any recognized divisions. Since there is large variability in the targeted 18S rRNA gene, amplification-based molecular methods can be problematic for eukaryotic organisms^92^. To address this issue some studies utilized a chloroplast 16S rRNA gene database for taxonomic assignments of photosynthetic eukaryotic organisms ^14^. For our study we sequenced the V9 region of 18S rRNA which has been shown to have a higher resolution at the genus level (80% identification rate) ^93^. However, genomic data from this part of BoB is very limited - therefore, the existing databases might have lower resolution in assigning the taxonomic profiles. Including other regions of the 18S rRNA, i.e., V2 and V4 might have recovered higher diversity of microbial eukaryotes in these regions.

The 16S rDNA based microbial profiling conducted in this study has revealed high bacterial diversity in the coastal regions of Cox’s Bazar and Saint Martin, In Cox’s Bazar, abundance of *Alteromonas*, *Methylophaga*, *Anaerospora*, *Marivita*, and *Vibrio* were identified, while in Saint Martin, the *Pseudoalteromonas*, *Nautella*, *Marinomonas*, *Vibrio*, and *Alteromonas* were dominating. A variety of factors, including the physical and chemical properties of the environment, the presence of other species, and human activities, can affect the composition of the bacterial community in a particular setting^94^. Therefore, variations in the physicochemical parameters may account for the disparities in bacterial dominance between Cox’s Bazar and Saint Martin. Overall, the surface aquatic community has been shown to be dominated by the Rhodobacteriaceae family, which are the major group of microorganisms involved in organic matter recycling in marine environments ^95^. The 16S rDNA-based metagenomic data analysis revealed the genus-level identification of *Alteromonas, Anaerospora, Methylophaga, Nautella, Marinomonas* and *Pseudoalteromonas* through the abundance of the OTUs 1, 5, 18, 2, 12 and 9 respectively in the samples S9, S11, S12, S7, S8 and S6. The Rhodobacteraceae family has been identified by 49 OTUs, a large number of which were classified to the genus level. The notable genera of Rhodobacteraceae are *Nautella, Anaerospora, Antarctobacter, Thalassobius, Thalassococcus, Roseivivax, and Roseovarius*. The Rhodobacteraceae family of bacteria typically flourish in marine settings and they mostly consist of aerobic photo- and chemoheterotrophs That are involved in symbiosis as well as contributors to sulfur and carbon biogeochemical cycles ^95^. The second most abundant family, the Flavobacteriaceae, have been identified by 45 different OTUs s, many of which were identified up to genus level. According to a previously published report, in the maritime environment, members of the bacterial family Flavobacteriaceae are extensively dispersed and frequently discovered in association with algae, fish, debris, or marine animals ^96,97^. The ability of marine Flavobacteriaceae to consume a variety of carbon sources is supported by the high frequency and diversity of genes encoding polymer-degrading enzymes, which are frequently organized in polysaccharide utilization loci (PULs) ^98,99^. With a high incidence of gene clusters encoding pathways for the generation of antibiotic, antioxidant, and cytotoxic chemicals, Flavobacteriaceae have a varied arsenal of secondary metabolite biosynthesis ^99^. Relatively higher abundance of the Flavobacteriaceae family in our study sites indicates the availability of complex macromolecules in these coastal regions.

From the sample-wide analysis of 16S data (Figure-4A), there were notable abundances of *Pseudoalteromonas, Alteromonas* and *Methylophaga* genus in S6, S9 and S12 respectively. *Pseudoalteromonas* is a recently recognized genus that includes many marine species that produce physiologically active compounds. Specifically, these species appear to produce several chemicals that have antimicrobial against a wide range of target organisms, which may benefit them in their competition for resources and surface colonization. *Pseudoalteromonas* species exhibit antibacterial, bacteriolytic, agarolytic, and algicidal properties and are typically found associated with marine eukaryotes ^100,101^. Additionally, several isolates of *Pseudoalteromonas* stop the growth of typical fouling species.

The genus *Alteromonas* have a wide range of habitats, including coastal and open ocean regions, deep sea and hydrothermal vents, and marine sediments ^102^. Since *Alteromonas* is known to have a wide variety of metabolic activities, including the breakdown of complex organic molecules ^103^. Among the other genera *Anaerospora*, *Marivita,* and *Vibrio* are also commonly found in marine environments, with *Vibrio* being of particular interest due to its several potentially pathogenic species ^104^. The presence of these bacteria in Cox’s Bazar water sample suggests that careful monitoring of their populations may be required to prevent potential negative impacts on human and animal health. The genus *Marinomonas*, which have been detected only in Jetty samples (S8), is considered as a promising candidate for potential biotechnological applications, such as the production of enzymes, biofuels, and biodegradable plastics ^105–107^.

Marine microorganisms exhibit numerous metabolic capabilities either as independent strains or as members of complex microbial consortia. They can produce eco-friendly chemicals and novel metabolites that can be used in the management and treatment of environmental waste, such as nontoxic biosurfactants and biopolymers and for the treatment of diseases ^108–111^. Many of the microbial lineages previously reported to synthesize antibiotic compounds have also been discovered in our study sites (Supplementary Data-3: Table-2). These include *Rhodobacteraceae bacterium* ^112^, *Pseudoalteromonas phenolica* ^113^, *Proteobacteria bacterium* ^114^, *Ruegeria* sp. ^115^, *Vibrio mediterranei* ^116^, *Phaeobacter* sp. ^117^ and *Marinomonas ostreistagni* ^118^ among others. Other microorganisms like *Alteromonas portus* ^119,120^ and *Seonamhaeicola algicola* ^121,122^ are known for production of antioxidants carotenoids, zeaxanthin; *Alteromonas oceani* ^123^ and *Ruegeria* sp. ^115^ for probiotics; *Alteromonas portus* ^120^ for anticancer activity; *Vibrio fortis* for biofouling ^124,125^ and *Phaeobacter italicus* for biodiesel prospects ^126,127^. Additionally, pathogens causing food borne illnesses like *Vibrio parahaemolyticus* have also been found.

Bangladesh has an extreme shortage of facilities and infrastructures for treatment of hospitals and municipal waste^128,129^. In fact, most wastes are disposed into the freshwater bodies, like rivers, canals, lakes etc., which eventually reach the estuarine and marine waters of the Bay of Bengal. This substantial agricultural runoff, as well as anthropogenic hospital and municipal discharge cause deposition of antibiotics and ARB in the surrounding coastal environment ^128^. Antimicrobial resistance (AMR) genes and residual antibiotics potentially impact the overall community composition and eventually threatening the ecological balance of microorganisms through unwanted exposure of autochthonous microbial community to the antimicrobial compounds and hereby disturbing the harmony of ecosystem health. It has already been documented that when naturally untainted environments are contaminated by ARB and ARGs, they can mobilize ARGs to naive bacterial communities ^130,131^. Although many studies have investigated the metabolic potential of the marine microbes in other oceanic regions, the functional and phylogenetic diversity of the microbial community in the coastal water of the BoB remain underexplored.

Our in-depth metagenomic analysis revealed presence of antibiotic resistance genes in multiple classes (Supplementary Data-3: Table 3 and 4) in the coastal microbial community of Saint-martin (S1) and Cox’s bazar (S2). Saint Martin Island microbial community harbored resistance genes against macrolides, aminoglycosides, and quinolones. On the other hand, the Cox’s bazar microbes contained larger spectrum of AMR genes, with higher coverage and abundance of each gene. These findings indicate the occurrence of antibiotic resistance genes in the surface waters of BoB, with higher abundance in the Cox’s Bazar region. As this area is highly inundated with tourists, all the year round, the coastal water encounters microbial populations originated from human and animals, allowing an intrusion and environmental adaptation of the allochthonous microbes into the natural microbial community. Besides, wastes from the coastal districts, including the second largest and populated city of Bangladesh “Chattogram”, are being dumped and carried away to the marine water through all the rivers connected to the BoB ^132–134^. Discharged waste coming from hospital and municipal sources contain reservoirs of antibiotics which are harbored in the feces of humans, chickens, and cows. Resistance against colistin-like last-resort antibiotics have been reported to be disseminated into the microbiome of marine water ^135^, although this was not found in the samples we studied. The resistomes of BoB microbiome strongly exemplifies how anthropogenic input can turn the coastal environment into a potential reservoir of antibiotic resistance, further threatening the public health. Given the implications for public health and marine ecological balance, future studies on the BoB coast as a potential sink and source of antibiotic resistance will be crucial.

The microbial profiling conducted in this study was produced using “universal” PCR primers, selected for their ability to simultaneously target both 16S and 18S rRNA genes. Microbial communities are now well understood as major contributors in maintaining balance in marine and terrestrial ecosystems. Despite being a highly dynamic tropical water body, the Indian Ocean has not attracted much attention from the scientists and remains the least explored source of its microbial biodiversity. Recent studies have added significantly through the use of metagenomics methods in marine microbial ecology. Ambient conditions shape microbiome responses to both short- and long-duration environment changes through processes including physiological acclimation, compositional shifts, and evolution.

Many open questions currently limit our capacity to assess how microbial processes influence the ecology of these environments, both under contemporary conditions and under future environmental change. Therefore, there is a clear need to prioritize and define key questions for future research that will allow for better assessments of how microbial processes truly influence the ecology and health of coastal marine environments.

## 5. Conclusion

The findings from this study provide the first insights into the properties, toxonomic composition and functional profiles of coastal microbial communities of the Bay of Bengal from Bangladesh. Our combined approach for 16S and 18S amplicon-based sequencing provides a much more comprehensive picture of the sublittoral epipelagic coastal water of BoB. The shotgun metagenomic analysis of these microbiomes reveals significantly abundant communities and their metabolic potential. The results could be potentially used in several downstream studies, such as the comparative analysis of coastal and deep-sea metagenomes to explore the bio-prospective potential of the Bay of Bengal.

## Data availability

The 16S, 18S and Shotgun sequences are available in **BioProject PRJNA936421, PRJNA936461** and **PRJNA936489**, respectively of NCBI database. All supplementary files are uploaded along with the manuscript.

## Acknowledgement

This study has received a generous instrument support from the Department of Microbiology, Jahangirnagar University. Reagents and consumable were provided partially grants from Ministry of Science and Technology, Govt. of Bangladesh, Faculty of Biological Sciences, Jahangirnagar University, Savar, Dhaka and Bangladesh, University Grants Commissions Bangladesh. There was no funding support for publication of the article.

## Authors contribution

SA and MSR performed bioinformatics analysis, visualized figures, interpreted results, and drafted the original manuscript. HA, KM, MMS, and FY carried out field experiments and curated the data. BM, SMG, NAH, and SRR edited and reviewed the manuscript. NAz, NAd, and SRR reviewed and edited the final draft, partial instrument supports were provided by SRR, partial reagent supports were provided by SA and MFA. MM and MFA conceived the study, availed the reagent support, critically reviewed the drafted manuscript, and supervised the research overall.

