## Supplementary data-3 for "Exploring Microbial Diversity and Functional Potential along the Bay of Bengal Coastline in Bangladesh: Insights from Amplicon Sequencing and Shotgun Metagenomics"

**Supplementary Data 2**

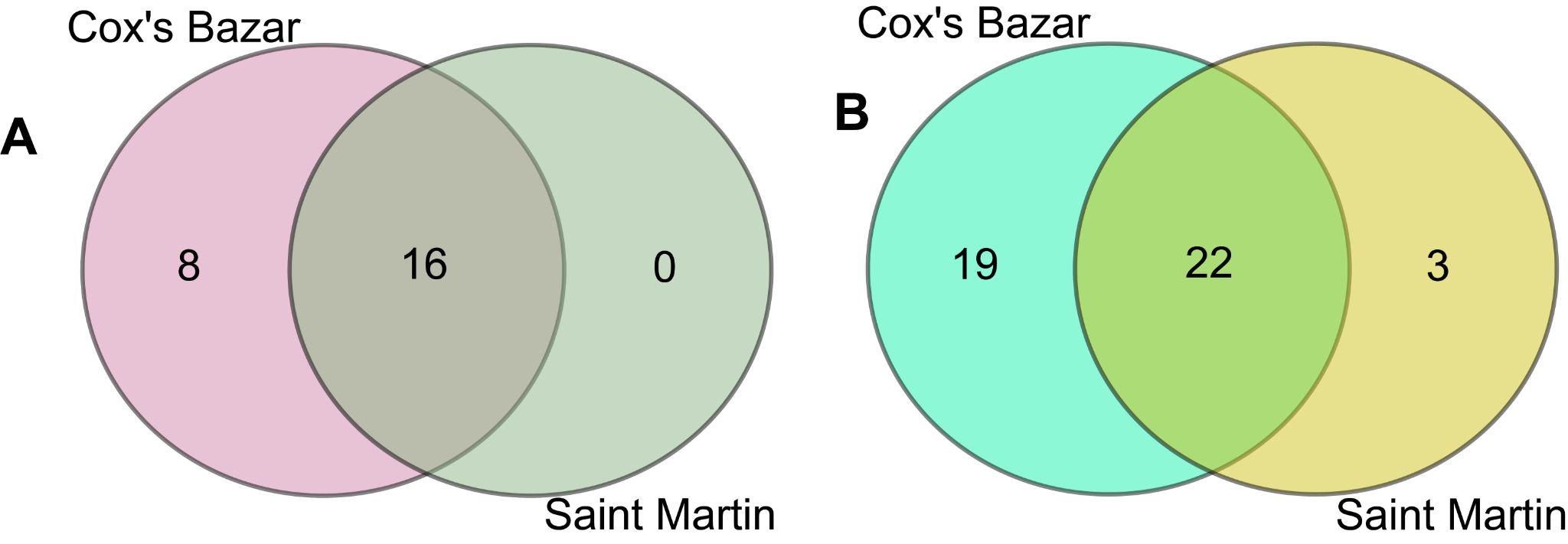
 **Supplementary Figure-1: Venn diagram of phylum and division level overlap of prokaryotic (A) and eukaryotic (B) samples from two locations.** Venn diagrams show common and unique microbial populations in the sampling locations.

**Figure-2: Comparison of relative abundance of forty-nine prokaryotic genus in the two different locations (Cox’s Bazar and Saint Martin).** The diversity for each genus is plotted on boxplots and comparisons are made with Wilcoxon sum rank test. Significance level (p-value) 0.0001, 0.001, 0.01, 0.05, and 0.1 are represented by the symbols "****", "***", "**", "*", and "n.s", respectively.

**SUPPLEMENTARY TABLES**

**Table-1: Presence of functional proteins based on BRITE Hierarchies level A, B and C in coastal marine water of BoB, Bangladesh.**

| **Level A** | **Level B** | **Level C** |
| --- | --- | --- |
| **Metabolism** | Carbohydrate metabolism | Glycolysis / Gluconeogenesis |
|  | Energy metabolism | Citrate cycle (TCA cycle ) |
|  | Lipid metabolism | Pentose phosphate pathway |
|  | Nucleotide metabolism | Pentose and glucuronate interconversions |
|  | Amino acid metabolism | Fructose and mannose metabolism |
|  | Metabolism of other amino acids | Galactose metabolism |
|  | Glycan biosynthesis and metabolism | Ascorbate and aldarate metabolism |
|  | Metabolism of cofactors and vitamins | Fatty acid biosynthesis |
|  | Metabolism of terpenoids and polyketides | Fatty acid elongation |
|  | Biosynthesis of other secondary metabolites | Fatty acid degradation |
|  | Xenobiotics biodegradation and metabolism | Cutin, suberine and wax biosynthesis |
|  | Not included in regular maps | Steroid biosynthesis |
| **Genetic Information Processing** | Transcription | Primary bile acid biosynthesis |
|  | Translation | Secondary bile acid biosynthesis |
|  | Folding, sorting and degradation | Ubiquinone and other terpenoid- quinone biosynthesis |
|  | Replication and repair | Steroid hormone biosynthesis |
|  | Information processing in viruses | Oxidative phosphorylation |
| **Environmental**  **Information Processing** | Membrane transport | Photosynthesis proteins |
|  | Signal transduction | Photosynthesis |
|  | Signaling molecules and interaction | Photosynthesis - antenna proteins |
| **Cellular Processes** | Transport and catabolism | Cytochrome P450 |
|  | Cell motility | Arginine biosynthesis |
|  | Cell growth and death | Purine metabolism |
|  | Cellular community - eukaryotes | Caffeine metabolism |
|  | Cellular community - prokaryotes | Pyrimidine metabolism |
| **Organismal Systems** | Aging | Alanine, aspartate and glutamate metabolism |
|  | Immune system | Tetracycline biosynthesis |
|  | Endocrine system | Aflatoxin biosynthesis |
|  | Circulatory system | Glycine, serine and threonine metabolism |
|  | Digestive system | Monobactam biosynthesis |
|  | Excretory system | Cysteine and methionine metabolism |
|  | Nervous system | Valine, leucine and isoleucine degradation |
|  | Sensory system | Geraniol degradation |
|  | Development and regeneration | Valine, leucine and isoleucine biosynthesis |
|  | Environmental adaptation | Lysine biosynthesis |
| **Human Diseases** | Cancer: overview | Lysine degradation |
|  | Cancer: specific types | Penicillin and cephalosporin biosynthesis |
|  | Immune disease | Arginine and proline metabolism |
|  | Neurodegenerative disease | Clavulanic acid biosynthesis |
|  | Substance dependence | Carbapenem biosynthesis |
|  | Cardiovascular disease | Prodigiosin biosynthesis |
|  | Endocrine and metabolic disease | Histidine metabolism |
|  | Infectious disease: bacterial | Tyrosine metabolism |
|  | Infectious disease: viral | Phenylalanine metabolism |
|  | Infectious disease: parasitic | Chlorocyclohexane and chlorobenzene degradation |
|  | Drug resistance: antimicrobial | Benzoate degradation |
|  | Drug resistance: antineoplastic | Bisphenol degradation |

**Table- 2: Metabolic functions of important bacterial species identified in the shotgun sequence of S1 and S2 samples from Bay of Bengal.**

| **Species** | **Functions** | **References** |
| --- | --- | --- |
| *Alteromonas* sp*.* | 1. Agarase-production  2. Metabolize aromatic hydrocarbons | 1. (Wang, Mou et al. 2006)  2. (Math, Jin et al. 2012) |
| *Rhodobacteraceae bacterium* | 1. Algicidal activity  2. Production of tropodithietic acid (TDA), a broad-spectrum antimicrobial compound | 1. (Zheng, Cui et al. 2015)  2. (Henriksen, Lindqvist et al. 2022) |
| *Pseudoalteromonas phenolica* | 1. Produces phenolic anti-MRSA substances  2. Recycle high-salt organic wastes | 1. (Isnansetyo and Kamei 2003)  2. (Song, Kim et al. 2020) |
| *Vibrio natriegens* | 1. High productivity in industrial fermentation  2. Biocatalyst for bioremediation of selenite | 1. (Hoffart, Grenz et al. 2017)  2. (Fernández-Llamosas, Castro et al. 2017) |
| *Alteromonas macleodii* | 1.Degrade multiple algal polysaccharides  2. Produce unusual high molecular-weight polysaccharide in the presence of glucose | 1. (Koch, Dürwald et al. 2019)  2. (Raguenes, Pignet et al. 1996) |
| *Proteobacteria bacterium* | 1. Ammonium-oxidation  2. Production of antimicrobial peptides | 1. (Voytek and Ward 1995)  2. (Desriac, Jégou et al. 2013) |
| *Marinomonas communis* | 1. Arsenic-accumulating bacteria  2.Ammonia and nitrite reducing organism | 1. (Takeuchi, Kawahata et al. 2007)  2. (Huang, Pan et al. 2020) |
| *Phaeobacter italicus* | 1.Improve yields of microalgae oils | 1. (Armstrong, Campos et al. 2022) |
| *Ruegeria sp.* | 1.Probiotics  2.Produces indigoidine, an antimicrobial agent | 1. (Rosado, Leite et al. 2019)  2. (Cude, Mooney et al. 2012) |
| *Vibrio sp.* | 1. Fermentation of sugars producing acids (formic, lactic, acetic, succinic acids), ethanol, and pyruvate. | 1. (Sampaio, Silva et al. 2022) |
| *Pseudoalteromonas sp.* | 1. Produce dimers and trimers from alginate.  2.Produce an extracellular serine protease which has algicidal activity. | 1. (Li, Dong et al. 2011)  2. (Lee, Kato et al. 2000) |
| *Marinomonas sp.* | 1. Produce silver nanoparticles (AgNPs) from silver nitrate. | 1. (John, Nagoth et al. 2020) |
| *Marisediminitalea aggregata* | 1. Biofilm formation on plastic surfaces. | 1. (Bos, Kaul et al. 2022) |
| *Vibrio mediterranei* | 1. Produce antibacterial compounds | 1. (Bruhn, Gram et al. 2007) |
| *Tritonibacter mobilis* | 1.Produce tropone derivative tropodithietic acid (TDA) an antimicrobial agent. | 1. (Henriksen, Lindqvist et al. 2022) |
| *Vibrio parahaemolyticus* | 1. Leading causes of seafood-borne illness | 1. (Davis, Jacobs et al. 2017) |
| *Alpha proteobacterium HIMB114* | 1.Transformation of organic carbon from labile to recalcitrant states. | 1. (Carini, Campbell et al. 2014) |
| *Candidatus Pelagibacter sp.* | 1.Degrade polycyclic aromatic hydrocarbons.  2. Nitrogen and carbon assimilation | 1. (Shi, Zhang et al. 2020)  2. (Partensky and Garczarek 2010) |
| *Alteromonas portus* | 1.Produce bioactive alginate oligosaccharides (ALO) with antioxidant and anticancer activity  2.Production of alginate lyase as antibiofilm agent | 1. (Jagtap, Sankar et al. 2022)  2. (Ethica, Zilda et al. 2021) |
| *Seonamhaeicola algicola* | 1.Saccharification of marine algae and fermentation of bioethanol  2. Source for the antioxidants | 1. (Zhou, Du et al. 2016)  2. (Brotosudarmo, Hardo et al. 2021) |
| *Alteromonas mediterranea* | 1.Degrade multiple algal polysaccharides | 1. (Koch, Dürwald et al. 2019) |
| *Phaeobacter sp.* | 1.Production of the Antimicrobial Secondary Metabolite Indigoidine | 1. (Cude, Mooney et al. 2012) |
| *Donghicola eburneus* | 1. Influence in steel corrosion | 1. (Moura, Ribeiro et al. 2018) |
| *Alteromonas oceani* | 1. High free radical scavenging ability  2. Probiotics | 1. (Dungan, Bulach et al. 2020) |
| *Vibrio fortis* | 1. Marine biofouling | 1. (Radjasa and Sabdono 2008) |
| *Oleibacter marinus* | 1. Degrades petroleum aliphatic hydrocarbons | 1. (Teramoto, Ohuchi et al. 2011) |
| *Ruegeria arenilitoris* | 1. Protects corals against pathogenic organisms | 1. (Miura, Motone et al. 2019) |
| *Marinomonas ostreistagni* | 1. Produce antifungal metabolites | 1. (Fields 2021) |
| *Pseudoalteromonas shioyasakiensis* | 1.Production of exopolysaccharides (EPSs) | 1. (Matsuyama, Sawazaki et al. 2014) |
| *Marinobacter sp.* | 1. Nitrate-assimilation  2. Capable of anaerobic sulfate and thiosulfate reduction. | 1. (Allen, Booth et al. 2005)  2. (Sigalevich, Baev et al. 2000) |
| *Candidatus Actinomarinales bacterium* | 1. Marine actinobacterial clade | 1. (López-Pérez, Haro-Moreno et al. 2020) |
| *Alteromonas alba* | 1. A novel species of the genus *Alteromonas* | 1. (Sun, Xamxidin et al. 2019) |
| *Vibrio alginolyticus* | 1. Production of thermolabile toxin | 1. (Sainz, Maeda-Martinez et al. 1998) |
| *Marivivens niveibacter* | 1. Novel spp. | 1. (Hu, Wang et al. 2018) |
| *Pseudoalteromonas agarivorans* | 1. Exopolysaccharide biosynthesis  2. Metalloprotease collagenase | 1. (Ju, Shan et al. 2022)  2. (Bhattacharya, Choudhury et al. 2018) |
| *Oleibacter sp.* | 1. Biosurfactant for contaminated soil and diesel degradation  2. Bioremediation of oil- contaminated water | 1. (Di Sia 2021)  2. (Catania, Lopresti et al. 2020) |
| *Shimia marina* | 1. [Exopolysaccharide](https://www.sciencedirect.com/topics/agricultural-and-biological-sciences/exopolysaccharide) and [secondary metabolite](https://www.sciencedirect.com/topics/agricultural-and-biological-sciences/secondary-metabolite) biosynthesis enzymes, degrade aromatic compounds. | 1. (Rodrigo-Torres, Pujalte et al. 2016) |
| *Alteromonas abrolhosensis* | 1. Xylose and glycerol assimilation | 1. (Nóbrega, Silva et al. 2018) |
| *Oceanospirillum sanctuarii* | 1. Contain *scv* locus, required for the production of polysaccharide | 1. (Sidhu, Thakur et al. 2017) |
| *Alteromonas marina* | 1. Agar degradation | 1. (Kolhatkar and Sambrani 2018) |
| *Alteromonas lipolytica* | 1.Poly-beta-hydroxybutyrate-producing bacterium | 1. (Shi, Wu et al. 2017) |
| *Vibrio maritimus* | 1. Siderophore-Producing Mutualistic Bacterium | 1. (Zhu, Cheng et al. 2022) |
| *Acidimicrobiaceae bacterium* | 1.Degradation of chlorinated ethenes tetra- and trichloro ethylene via oxygenase. | 1. (Ge, Huang et al. 2019) |
| *SAR86 cluster bacterium* | 1.Degradation of dimethyl sulfoniopropionate (DSMP) and cloud formation | 1. (González, Simó et al. 2000) |
| *Pseudoalteromonas spongiae* | 1. Larval settlement | 1. (Huang, Dobretsov et al. 2007) |
| *Halieaceae bacterium* | 1. Steroid degradation | 1. (Holert, Cardenas et al. 2018) |
| *Phaeobacter inhibens* | 1. Large number of Amino Acid and Sugar Catabolism | 1. (Wünsch, Trautwein et al. 2019) |
| *Rhodobacterales bacterium* | 1. Biocatalyst of glycidyl phenyl ether | 1. (Woo, Kang et al. 2010) |

**Table 3:** Antimicrobial resistance gene profiling for S1 (Saint martin) sample.

| **Gene Family/ Group** | **Type** | **Mechanisms** | **Class** | **% Gene Coverage** | **% Of identical matches** | **Abundance** |
| --- | --- | --- | --- | --- | --- | --- |
| **Mapped reads:** NDARO database with EzBioCloud pipeline (mapped with the gene by bowtie2 with the --very-sensitive option) | | | | | | |
| qnrVC | Drug | quinolone resistance pentapeptide repeat protein QnrVC1 | Quinolone | 99.39 | ≥80 | 2265 |
| dfrA6 | Drug | trimethoprim-resistant dihydrofolate reductase DfrA6 | Trimethoprim | 92.19 | ≥80 | 1780 |
| qnrVC | Drug | quinolone resistance pentapeptide repeat protein QnrVC4 | Quinolone | 87.06 | ≥80 | 2547 |
| **Mapped Reads:** MEGARes database with AMR++ Pipeline (BWA and resistome analyzer) | | | | | | |
| CAP16S group | Drugs | Cationic peptide-resistant 16S ribosomal subunit protein | Cationic antimicrobial peptides | 90.92 | ≥80 | 554 |
| A16S group | Drugs | Aminoglycoside-resistant 16S ribosomal subunit protein | Aminoglycosides | 99 | ≥80 | 3111 |
| *erm* | Drugs | Macrolide-resistant 23S rRNA mutation | MLS | 95 | ≥80 | 7339 |
| *rrs*C | Drugs | Aminoglycoside-resistant 16S ribosomal subunit protein | Aminoglycosides | 88.2 | ≥80 | 550 |
| *rrs*H | Drugs | Aminoglycoside-resistant 16S ribosomal subunit protein | Aminoglycosides | 89.43 | ≥80 | 621 |
| *rps*L |  | Aminoglycoside-resistant 16S ribosomal subunit protein | Aminoglycosides | 84.53 | ≥80 | 89 |
| TUFAB group | Drugs | EF-Tu inhibition | Elfamycins | 99 | ≥80 | 1359 |
| **Assembled data (Contigs;** MEGARes database with BLAST pipeline**)** | | | | | | |
| *vme*Z | Biocides | Multi-biocide RND efflux pump | Multi-biocide resistance | 99.84 | 81.71 | NA |
| *vme*D | Biocides | Multi-biocide RND efflux pump | Multi-biocide resistance | 99.78 | 80.66 | NA |
| TUFAB group | Drugs | EF-Tu inhibition | Elfamycins | 99.49 | 88.81 | NA |
| *gyr*A | Drugs | Fluoroquinolone-resistant DNA topoisomerases | Fluoroquinolones | 78.70 | 83.02 | NA |

**Table 4:** Antimicrobial resistance gene profiling for S2 sample.

| **Gene Family/ Group** | **Type** | **Mechanisms** | **class** | **% Gene Coverage** | **% Of identical matches** | **Abundance** |
| --- | --- | --- | --- | --- | --- | --- |
| **Raw reads mapped:** NDARO database with EzBioCloud pipeline (mapped with the gene by bowtie2 with the --very-sensitive option) | | | | | | |
| *tet(G)* | AMR | tetracycline efflux MFS transporter Tet(G) | Tetracycline | 100.00 | ≥80 | 238355 |
| *ere(A)* | AMR | EreA family erythromycin esterase | Macrolide | 100.00 | ≥80 | 18865 |
| *qnrS* | AMR | quinolone resistance pentapeptide repeat protein QnrS2 | Quinolone | 100.00 | ≥80 | 131607 |
| *floR2* | AMR | chloramphenicol/florfenicol efflux MFS transporter FloR2 | Phenicol | 100.00 | ≥80 | 345132 |
| *mph(F)* | AMR | Mph(F) family macrolide 2'-phosphotransferase | Macrolide | 100.00 | ≥80 | 9773 |
| *qacE* | BIOCIDE | quaternary ammonium compound efflux SMR transporter QacE delta 1 | Quaternary ammonium | 100.00 | ≥80 | 6566 |
| *tet(X)* | AMR | tetracycline-inactivating monooxygenase Tet(X) | Tetracycline | 98.71 | ≥80 | 9946 |
| *erm(F)* | AMR | 23S rRNA (adenine(2058)-N(6))-methyltransferase Erm(F) | Macrolide | 96.63 | ≥80 | 4358 |
| *tet(G)* | AMR | tetracycline efflux MFS transporter Tet(G) | Tetracycline | 94.56 | ≥80 | 29142 |
| *sul2* | AMR | sulfonamide-resistant dihydropteroate synthase Sul2 | Sulfonamide | 92.43 | ≥80 | 7190 |
| *aadA1* | AMR | ANT(3'')-Ia family aminoglycoside nucleotidyltransferase AadA1 | Aminoglycoside | 91.29 | ≥80 | 4074 |
| *aac(6')-Ib11* | AMR | aminoglycoside N-acetyltransferase AAC(6')-Ib11 | Aminoglycoside | 90.35 | ≥80 | 1637 |
| *sul1* | AMR | sulfonamide-resistant dihydropteroate synthase Sul1 | Sulfonamide | 90.25 | ≥80 | 5134 |
| *ere(D)* | AMR | EreD family erythromycin esterase | Macrolide | 90.22 | ≥80 | 5770 |
| *sul2* | AMR | sulfonamide-resistant dihydropteroate synthase Sul2 | Sulfonamide | 88.11 | ≥80 | 3922 |
| *aac(6')-Ib* | AMR | AAC(6')-Ib family aminoglycoside 6'-N-acetyltransferase | Aminoglycoside | 86.85 | ≥80 | 2710 |
| *sul2* | AMR | sulfonamide-resistant dihydropteroate synthase Sul2 | Sulfonamide | 84.51 | ≥80 | 6743 |
| *qacE* | BIOCIDE | quaternary ammonium compound efflux SMR transporter QacE | Quaternary ammonium | 82.58 | ≥80 | 1057 |
| *aac(6')-Ib* | AMR | AAC(6')-Ib family aminoglycoside 6'-N-acetyltransferase | Aminoglycoside | 80.54 | ≥80 | 1631 |
| ***Raw reads mapped:*** *MEGARes database with AMR++ Pipeline (BWA and resistome analyzer)* | | | | | | |
| *A16S group* | AMR | Aminoglycoside-resistant_16S_ribosomal_subunit_protein | Aminoglycosides | 99.00 | ≥80 | 2703 |
| *CAP16S group* | AMR | Cationic_peptide-resistant_16S_ribosomal_subunit_protein | Cationic_antimicrobial_peptides | 94.88 | ≥80 | 539 |
| *catB* | AMR | Chloramphenicol_acetyltransferases | Phenicol | 88.90 | ≥80 | 18 |
| *floR* | AMR | Phenicol_resistance_MFS_efflux_pumps | Phenicol | 99.91 | ≥80 | 500 |
| *merC* | Metals | Mercury_resistance_protein | Mercury_resistance | 99.76 | ≥80 | 21 |
| *merT* | Metals | Mercury_resistance_protein | Mercury_resistance | 99.72 | ≥80 | 16 |
| *MLS23S group* | Drugs | Macrolide-resistant_23S_rRNA_mutation | MLS | 90.00 | ≥80 | 5929 |
| *mphE* | Drugs | Macrolide_phosphotransferases | MLS | 99.89 | ≥80 | 22 |
| *QACEDELTA1* | Multi-compound | Drug_and_biocide_SMR_efflux_pumps | Drug_and_biocide_resistance | 82.81 | ≥80 | 2 |
| *qnrS* | Drugs | Quinolone_resistance_protein_Qnr | Fluoroquinolones | 99.85 | ≥80 | 178 |
| *qnrS* | Drugs | Quinolone_resistance_protein_Qnr | Fluoroquinolones | 96.66 | ≥80 | 92 |
| *rrsC* | Drugs | Aminoglycoside-resistant_16S_ribosomal_subunit_protein | Aminoglycosides | 93.71 | ≥80 | 544 |
| *rrsH* | Drugs | Aminoglycoside-resistant_16S_ribosomal_subunit_protein | Aminoglycosides | 95.01 | ≥80 | 530 |
| *tetG* | Drugs | Tetracycline_resistance_MFS_efflux_pumps | Tetracyclines | 99.90 | ≥80 | 694 |
| *TUFAB group* | Drugs | EF-Tu_inhibition | Elfamycins | 94.01 | ≥80 | 884 |
| ***Assembled data (Contigs)*** *MEGARes database with BLAST pipeline* | | | | | | |
| *floR* | Drugs | Phenicol_resistance_MFS_efflux_pumps | Phenicol | 99.92 | 99.92 | NA |
| *A16S group* | Drugs | Aminoglycoside-resistant_16S_ribosomal_subunit_protein | Aminoglycosides | 99.87 | 87.74 | NA |
| *qnrS* | Drugs | Quinolone_resistance_protein_Qnr | Fluoroquinolones | 99.85 | 99.54 | NA |
| *TUFAB group* | Drugs | EF-Tu_inhibition | Elfamycins | 98.65 | 83.99 | NA |
| *merR1* | Metals | Mercury_resistance_regulator | Mercury_resistance | 89.89 | 95.41 | NA |

**Table 3:** Virulence factors genes in S1 sample.

| **Name** | **Category** | **VFGID** | **% Gene Coverage** | | **% Of identical matches** | **Abundance (read counts)** | **Database and pipeline** |
| --- | --- | --- | --- | --- | --- | --- | --- |
| **Mapped reads** | | | | | | | |
| Flagellar basal body protein (*flg*B) | Motility;Flagella-mediated motility | VFG007360 | 100 | ≥80 | | 7376 | VFDB database with EzBioCloud pipeline (mapped with the gene by bowtie2 with the --very-sensitive option) |
| Flagellar basal body rod protein (*flg*C) | Motility;Flagella-mediated motility | VFG007354 | 99.75 | ≥80 | | 6317 |  |
| Flagellar basal body rod modification protein (*flg*D) | Motility;Flagella-mediated motility | VFG007348 | 85.73 | ≥80 | | 3319 |  |
| Putative MSHA pilin protein MshA (*msh*A) | Adherence;Fimbrial adhesin;Type IV pili;Type IVa pili (T4aP) | VFG006988 | 84.77 | ≥80 | | 1812 |  |
| Flagellar biosynthesis sigma factor FliA (*fli*A) | Motility;Flagella-mediated motility | VFG007576 | 83.26 | ≥80 | | 3020 |  |
| Hypothetical protein (*pvs*B) | Nutritional/Metabolic factor;Metal uptake;Iron Uptake | VFG044180 | 80.68 | ≥80 | | 2396 |  |
| Polar flagellar rod protein FlaI (*fla*I) | Motility;Flagella-mediated motility | VFG007510 | 79.74 | ≥80 | | 302 |  |
| Putative diaminopimelate decarboxylase protein (*pvs*E) | Nutritional/Metabolic factor;Metal uptake;Iron Uptake | VFG044183 | 78.39 | ≥80 | | 3920 |  |
| Putative ferrichrome ABC transporter permease (*pvu*C) | Nutritional/Metabolic factor;Metal uptake;Iron Uptake | VFG044175 | 77.07 | ≥80 | | 2860 |  |
| Putative transport protein (*pvs*C) | Nutritional/Metabolic factor;Metal uptake;Iron Uptake | VFG044181 | 74.63 | ≥80 | | 2393 |  |
| Chemotaxis protein CheY (*che*Y) | Motility;Flagella-mediated motility | VFG007570 | 74.54 | ≥80 | | 2547 |  |
| **Mapped reads** | | | | | | | |
| Chemotaxis protein CheY (*che*Y) | Motility;Flagella-mediated motility | VFG007570 | 100.00 | ≥80 | | 29 | VFDB database with AMR++ Pipeline (BWA and resistome analyzer) |
| Flagellar basal body rod protein FlgB (*flg*B) | Motility;Flagella-mediated motility | VFG007360 | 100.00 | ≥80 | | 27 |  |
| Flagellar basal-body rod protein FlgF (*flg*F) | Motility;Flagella-mediated motility | VFG007336 | 92.27 | ≥80 | | 25 |  |
| Flagellar basal-body rod modification protein FlgD (*flg*D) | Motility;Flagella-mediated motility | VFG007348 | 99.72 | ≥80 | | 24 |  |
| Flagellin (*fla*B) | Motility;Flagella-mediated motility | VFG007528 | 92.75 | ≥80 | | 24 |  |
| Flagellar biosynthesis sigma factor FliA (*fli*A) | Motility;Flagella-mediated motility | VFG007576 | 92.24 | ≥80 | | 19 |  |
| Type II secretion system major pseudopilin GspG (*eps*G) | Effector delivery system | VFG007086 | 91.89 | ≥80 | | 18 |  |
| Flagellar rod protein FlaI (*fla*I) | Motility;Flagella-mediated motility | VFG007510 | 93.79 | ≥80 | | 16 |  |
| Flagellar basal body rod protein FlgC (*flg*C) | Motility;Flagella-mediated motility | VFG007354 | 100.00 | ≥80 | | 13 |  |
| (mshA) MSHA pilin protein MshA | Adherence | VFG006988 | 88.89 | ≥80 | | 8 |  |
| **Assembly based (Contigs)** | | | | | | | |
| Flagellar basal-body rod protein FlgF(*flg*F) | Motility;Flagella-mediated Motility | VFG007336 | 99.87 | 88.00 | | NA | VFDB database with BLAST pipeline |
| Flagellar basal body rod protein FlgC (*flg*C) | Motility;Flagella-mediated Motility | VFG007354 | 99.76 | 92.75 | | NA |  |
| Type II secretion system minor pseudopilin GspI (*gsp*I) | Effector delivery system | VFG007074 | 99.49 | 83.81 | | NA |  |
| MSHA pilin protein MshA (*msh*A) | Adherence | VFG006988 | 91.03 | 85.51 | | NA |  |
| MSHA biogenesis protein MshH (*msh*H) | Adherence | VFG006922 | 82.84 | 80.53 | | NA |  |
| Flagellar biosynthesis sigma factor FliA (*fli*A) | Motility;Flagella-mediated Motility | VFG007576 | 79.86 | 85.91 | | NA |  |
| **Assembly based (Bins)** | | | | | | | |
| MSHA biogenesis protein MshH (*msh*H) | Adherence | VFG006922 | 99.95 | 80.09 | | NA | VFDB database with BLAST pipeline |
| Flagellar hook-associated protein 1 FlgK (*flg*K) | Motility;Flagella-mediated motility | VFG007306 | 99.95 | 82.74 | | NA |  |
| Type II secretion system ATPase GspE (*eps*E) | Effector delivery system | VFG007098 | 99.93 | 82.46 | | NA |  |
| MSHA biogenesis protein MshG (*msh*G) | Adherence | VFG006970 | 99.92 | 80.77 | | NA |  |
| Sodium-type flagellar protein MotY (*mot*Y) | Motility;Flagella-mediated Motility | VFG007612 | 99.89 | 80.18 | | NA |  |
| MSHA biogenesis protein MshM (*msh*M) | Adherence | VFG006952 | 99.88 | 80.38 | | NA |  |
| Flagellar motor protein (*mot*A) | Motility;Flagella-mediated motility | VFG007600 | 99.87 | 81.50 | | NA |  |
| Flagellar biosynthesis protein (flhA) | Motility;Flagella-mediated motility | VFG007594 | 99.86 | 82.80 | | NA |  |
| Sodium-type flagellar protein MotX (*mot*X) | Motility;Flagella-mediated motility | VFG007618 | 87.11 | 82.34 | | NA |  |
| Twitching motility protein PilT (*pil*T) | Adherence | VFG013907 | 83.86 | 85.73 | | NA |  |
